# Molecular models for Gram-positive bacterial strains: Assessing membrane properties and small molecule interactions for *S. aureus*, *S. epidermidis* and *N. lacusekhoensis*

**DOI:** 10.64898/2026.07.10.737677

**Authors:** Rakesh Vaiwala, Ernest Christy, Morris Waskar, K. Ganapathy Ayappa

## Abstract

We present a comparative study of the inner membrane of three Gram-positive bacterial strains, namely *S. aureus*, *S. epidermidis* and *N. lacusekhoensis*. A lipidomics study is used to obtain the lipid architecture and composition for *S. epidermidis* found in the skin microbiome and *N. lacusekhoensis*, an extremophile present in halophilic and alkophilic environments. Differences between the strains arise from both the lipid architecture and the cardiolipin content varying from 5% in *S. aureus* to 85% in *N. lacusekhoensis*. We develop coarse grained (CG) Martini-3 membrane models which reproduce structural properties such as membrane area, thickness, density distributions as well as ion-correlations with all-atom models. Inter-lipid correlations reveal a homogeneous distribution of lipids in the membranes despite the wide variation in lipid types and composition. Mechanical properties such as the area stretch modulus increased with cardiolipin content, however the bending modulus has a more complex dependence on membrane charge and lipid type. Using the CG models we evaluate the insertion free energies for four widely used antimicrobial molecules. Entry barriers for thymol and methylparaben arise from the charge density modulation at the membrane headgroups due to counterion condensation. The entry mechanisms of the antimicrobial peptide cecropin-melittin-15 (CM15) and the preservative molecule ethyl-lauroyl-arginate (ELAR) are found to be similar across all three strains. We also illustrate the manner in which the extremophilic strain, *N. lacusekhoensis* with its high cardiolipin content, modulates the partitioning kinetics of the antimicrobial molecule thymol with pH and salt. Our study reveals that membrane properties are largely conserved across the three model membranes. The molecular models and insights emerging from the present work should aid in the development of novel antimicrobials against Gram-positive strains.

## 1 INTRODUCTION

Lipid diversity and compositional variations in the bacterial cell envelope dictate membrane properties for bacterial species and modulate the transport of external molecules. ^1,2^ The lipid chemistry and architecture in Gram-positive bacteria differ from those found in Gram-negative bacteria. ^3^ Gram-positive strains such as *Staphylococcus aureus* (*S. aureus*) and *Staphylococcus epidermidis* (*S. epidermidis*), which inhabit the human skin microbiota, are usually considered commensal bacteria, protecting the skin against pathogenic strains. ^4,5^ *Nesterenkoina lacusekhoensis* (*N. lacusekhoensis*) belongs to another class of Gram-positive extremophile strains that survive and grow in high salt and alkaline conditions. It is found in sea water in both spherical (cocci) and rod shaped (bacilli) morphologies. ^6^ To survive and withstand the osmotic pressure gradient caused by high water salinity in the extracellular environment, bacteria accumulate organic solutes (osmolytes) in their cytoplasm and thereby balance the osmotic stresses. ^7,8^ A key factor that contributes to the survival of bacteria in extreme environments is the ability to modulate lipid chemistry using appropriate lipid synthesis pathways. *S. aureus* is perhaps the most widely studied Gram-positive strain and a few molecular dynamics (MD) simulations have been reported on the inner phospholipid membrane. ^9–14^ However *S. epidermidis* and *N. lacusekhoensis* have not been investigated using MD simulations as the lipid composition is yet to be reported in the literature. In order to build both atomistic and coarse-grained models we carried out a lipidomics study to determine the lipid composition for these strains. Using the lipidomics data we developed molecular models in the all-atom (AA) and coarse grained (CG) descriptions, and carried out MD simulations to compare and contrast the influence of lipid composition on membrane structure and biophysical properties for three Gram-positive bacterial strains, namely *S. aureus*, *S. epidermidis* and *N. lacusekhoensis*.

The cytoplasmic membrane lipids for Gram-positive strains contain fatty acids with varying lipid saturation and branching that occurs either on one or two carbons from the terminal carbon of the lipid chain. These are referred to as *iso* and *anteiso* branching, respectively, and they are known to modulate the membrane properties. ^15,16^ Molecular simulations have shown that branching increases membrane fluidity and chain disorder. ^17^ The branching lowers melting transition to facilitate survival of bacteria at lower temperatures. ^18^ Furthermore, the phospholipid headgroups play a major role in modulating membrane functions. Apart from having the phosphatidyl glycerol headgroups, membrane lipids are also modified to include amino acid residues such as lysine in *S. aureus* and glycerol in *S. edipermidis*. Another important and lesser studied component in bacterial membranes is the four tailed cardiolipin (diphosphatidylglycerol), which is a prominent component in mitochondrial membranes. ^19^ Although bacterial inner membranes have typically 5-10% of cardiolipin, cardiolipin content can be as high as 65-80% in extremophile strains, allowing them to survive in high salt conditions. To our knowledge there is no systematic study of the influence of cardiolipin content on Gram-positive bacterial membrane properties. A few MD simulations have been devoted to Gram-positive *S. aureus* strains where the cardiolipin content is typically ∼ 5%. ^9–11,14^

A recent simulation work on *S. aureus* membrane incorporated multicomponent lipids with varying range of acyl chains containing *iso* and *anteiso* branching, and the study established an impact of branched lipids on lipid ordering and membrane structure. ^12^ Coarse-grained MD simulations have also investigated the influence of the number of hydrocarbon chains in cardiolipin on the bulk phase behavior. ^20,21^ Cardiolipin has found to alter ion binding, water dipole orientations, lipid diffusion, hydrocarbon chain order and a propensity to bind to inner membrane proteins. ^21–24^

The atomic insights emerging from MD simulations depends on the appropriate choice of molecular models and force fields. On account of the large diversity of lipid types and compositions, molecular models and interpretations pertaining to one strain of bacteria cannot be directly translated and applied to other bacterial strains. MD simulations have predominantly been carried out on Gram-negative bacteria, e.g. *Escherichia coli*, while the simulation studies on Gram-positive strains are restricted mostly to *S. aureus* membranes. In this work we have developed both all-atom (AA) and coarse-grained (CG) models for three Gram-positive strains with distinctly varying membrane compositions. In particular, the cardiolipin content ranges from low (5%), moderate (22%) to high (85%) in *S. aureus*, *S. epidermidis* and *N. lacusekhoensis* strains, respectively. The membrane lipid compositions for *S. epidermidis* and *N. lacusekhoensis* were obtained from independent lipidomics data analysis carried out as a part of this study. The CG models for these three strains are developed within the Martini-3 framework, and extensive membrane simulations are carried out to validate the CG models with the atomistic simulations using CHARMM36 force field. Using the CG models, we compared properties across the three Gram-positive strains. These include charge densities, mechanical properties and free energy of insertion of small molecule antimicrobials as well as the antimicrobial peptide, CM15. In addition, we carried out atomistic simulations to explore the influence of salt and pH on insertion free energies of thymol on *N. lacusekhoensis* to illustrate the interplay between salt and pH that confer the halophilic properties to these Gram-positive strains.

## 2 LIPIDOMICS: MATERIALS AND METHODS

### 2.1 Bacterial Growth Media

*S. epidermidis* ATCC 12228 was purchased from ATCC, while *N. lacusekhoensis* was isolated from local marketed soaps and identified by 16S rRNA sequencing. *S. epidermidis* was maintained on Tryptic Soy Agar (TSA) and Tryptic Soy Broth (TSB). *N. lacusekhoensis* was maintained on modified TSA plates containing 1.5% NaCl and 1% of 70 Total Fatty Matter (TFM) bar soap powder.

### 2.2 Bacterial Lipidomics

*S. epidermidis* ATCC 12228 was cultured in TSB for 24 hours at 37 ^◦^C and collected by centrifugation at 5000 g for 5 minutes. The resulting pellets were washed three times with sterile distilled water and then adjusted to an optical density (OD) of 1 at 600 nm. In the case of *N. lacusekhoensis* (environmental isolate), bacteria were grown on modified TSA plates for 7 days at 30-32^◦^C. The culture was gently scraped, collected, washed, and OD was adjusted as described for *S. epidermidis*. Next, 10 mL of this suspension was centrifuged, the supernatant was removed, the pellet was quickly frozen in liquid nitrogen and stored at 80^◦^C until the cell lysis step. For cell lysis, samples were thawed to 20-25^◦^C, resuspended in 0.5 mL distilled water, transferred to 2 mL Eppendorf tubes, and cooled over ice bath. Glass beads (0.5 mm) were added to the suspensions, and cells were disrupted using a Mini-bead beater (1 minute run and 1 minute rest, repeated for 5 cycles). Samples were kept on ice for 5 minutes, after which the supernatant was transferred to fresh tubes and stored at 80^◦^C until shipment. Finally, samples were shipped in dry ice to Lipotype GmbH, Germany, for lipid analysis.

Lipid analysis of samples was carried out by Lipotype GmbH, Germany, using their proprietary Shotgun Lipidomics platform. Briefly, it consists of automated sample extraction, automated direct infusions, and high-resolution Orbitrap mass spectrometry including lipid class-specific internal standards to assure absolute quantification of lipids. Lipid identification from mass spectra was done using their in-house developed software - LipotypeXplorer, and analyses were performed using the Lipotype laboratory information and management system and LipotypeZoom - a web browser-based data visualization tool. Samples were spiked with lipid class-specific internal standards and were then extracted using HPLC/LC-MS analytical grade chloroform and methanol. ^25^ After drying and re-suspending in MS acquisition mixture, lipid extracts were subjected to mass spectrometric analysis. Mass spectra were acquired on a hybrid quadrupole/Orbitrap mass spectrometer equipped with an automated nano-flow electrospray ion source in both positive and negative ion mode. Lipid identification using LipotypeXplorer was performed on unprocessed mass spectra. ^26^ For MS-only mode, lipid identification was based on the molecular masses of the intact molecules. MSMS mode included the collision-induced fragmentation of lipid molecules and lipid identification was based on both the intact masses and the masses of the fragments. The dynamic range for samples was determined prior to analysis. ^25^ Based on these data, limits of quantification and coefficients of variation for the different lipid classes were determined. Limits of quantification were in the lower *µ*M to sub-*µ*M range, depending on the lipid class. The average coefficient of variation for a complete set of quantified lipid classes was around 10-15%. Each analysis was accompanied by a set of blank samples to control for a background and a set of quality control reference for *S. epidermidis* and *N. lacusekhoensis* samples to control for intra-run reproducibility and sample specific issues.

Shotgun lipidomics revealed distinct membrane lipid composition profiles for *S. epidermidis* ATCC 12228 and *N. lacusekhoensis*. Figure 1 summarizes the relative abundances of major lipid classes detected in the two species. These profiles illustrate clear species specific differences in lipid architecture, reflecting the unique membrane adaptations of each organism. For *N. lacusekhoensis*, fatty acid profiling showed that saturated and branched chain fatty acids dominated the lipidome. The three most abundant components were FA 16:0 (32.17%), FA 18:1 (23.50%), and FA 15:0 (13.82%). Additional notable contributors included FA 16:0 iso (11.69%) and FA 17:0 (9.16%), with the remaining fatty acids collectively representing 9.66% of the total composition, as given in Table S1 in Supporting Information (SI). In contrast, *S. epidermidis* displayed a different acyl chain distribution pattern, characterized by a mixture of *iso* and *anteiso* branched saturated chains. Key contributors included FA 14:0 iso, FA 15:0 iso, FA 15:0 anteiso, FA 18:0, and FA 20:0, with minor components grouped under other chains. These variations underscore differences in membrane fluidity and structural organization between the two species. Overall, the lipidomic profiles highlight fundamental biochemical distinctions between *S. epidermidis* and *N. lacusekhoensis*, reflecting their ecological niches and physiological adaptations.

**Figure 1:**
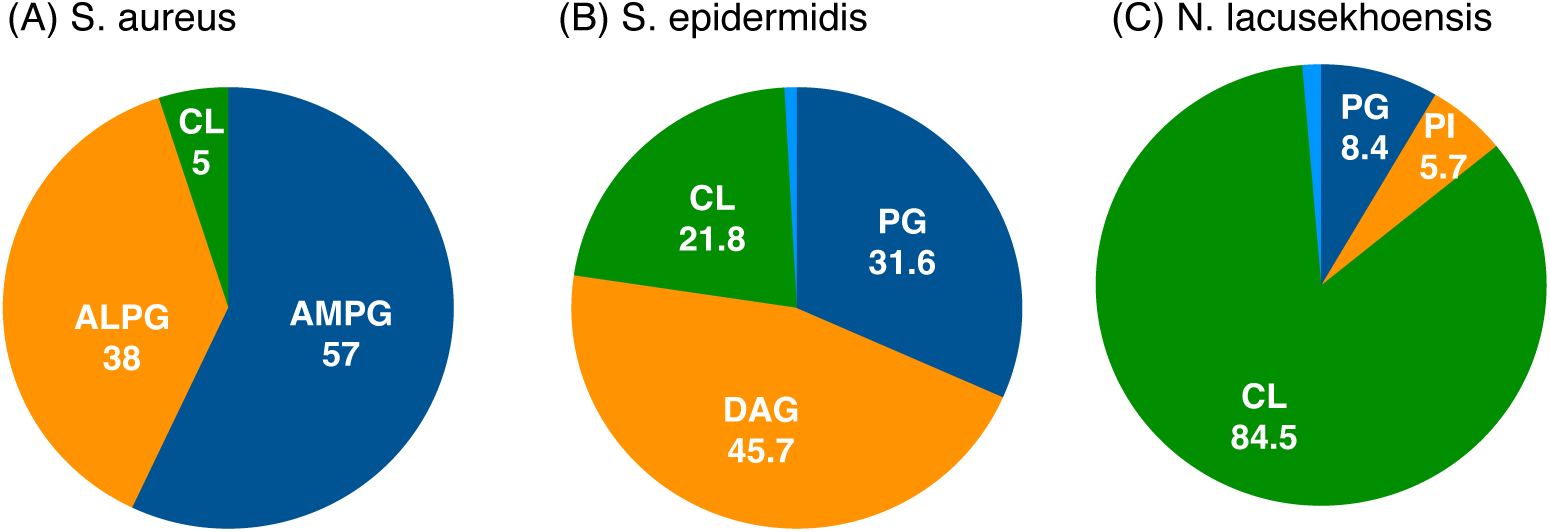
Lipid distribution profiles depicting relative abundances of major lipid classes for (A) *S. aureus* (B) *S. epidermidis*, and (B) *N. lacusekhoensis*. PG, phosphatidylglycerol; LPG, lysyl-phosphatidylglycerol; DAG, diacylglycerol; CL, cardiolipin; PI, phosphatidylinositol. The minor fractions (blue) are other lipids.

## 3 LIPID COMPOSITIONS AND MOLECULAR MODELS

The cytoplasmic membrane of *S. aureus* is composed of 57% phosphatidylglycerol (PG), 38% Lysyl-PG and 5% cardiolipin. ^27,28^ Our lipidomics data on *S. epidermidis* (ATCC 12228) and *N. lacusekhoensis* reveal a detailed compositions for membrane lipids, with a wider range of lipid chain length, carbon unsaturation, asymmetry in acyl chains, anteiso-branching in lipid tails, and a variety of headgroups composed of diacylglycerol (DAG), phosphatidylglycerol (PG), lysine (lysyl-PG), diphosphatidylglycerol (cardiolipin), and phosphatidylinositol (PI) moieties. We have considered the lipids with major compositions to construct the membrane models in this study (Table 1). The cardiolipin content in *N. lacusekhoensis* is 85%, while the membrane of *S. epidermidis* contains 46% of DAG molecules and 22% of cardiolipin. The lipid structures are shown in Figure 2, which also depicts the CG beads and the mapping schemes in the framework of Martini-3.

**Figure 2:**
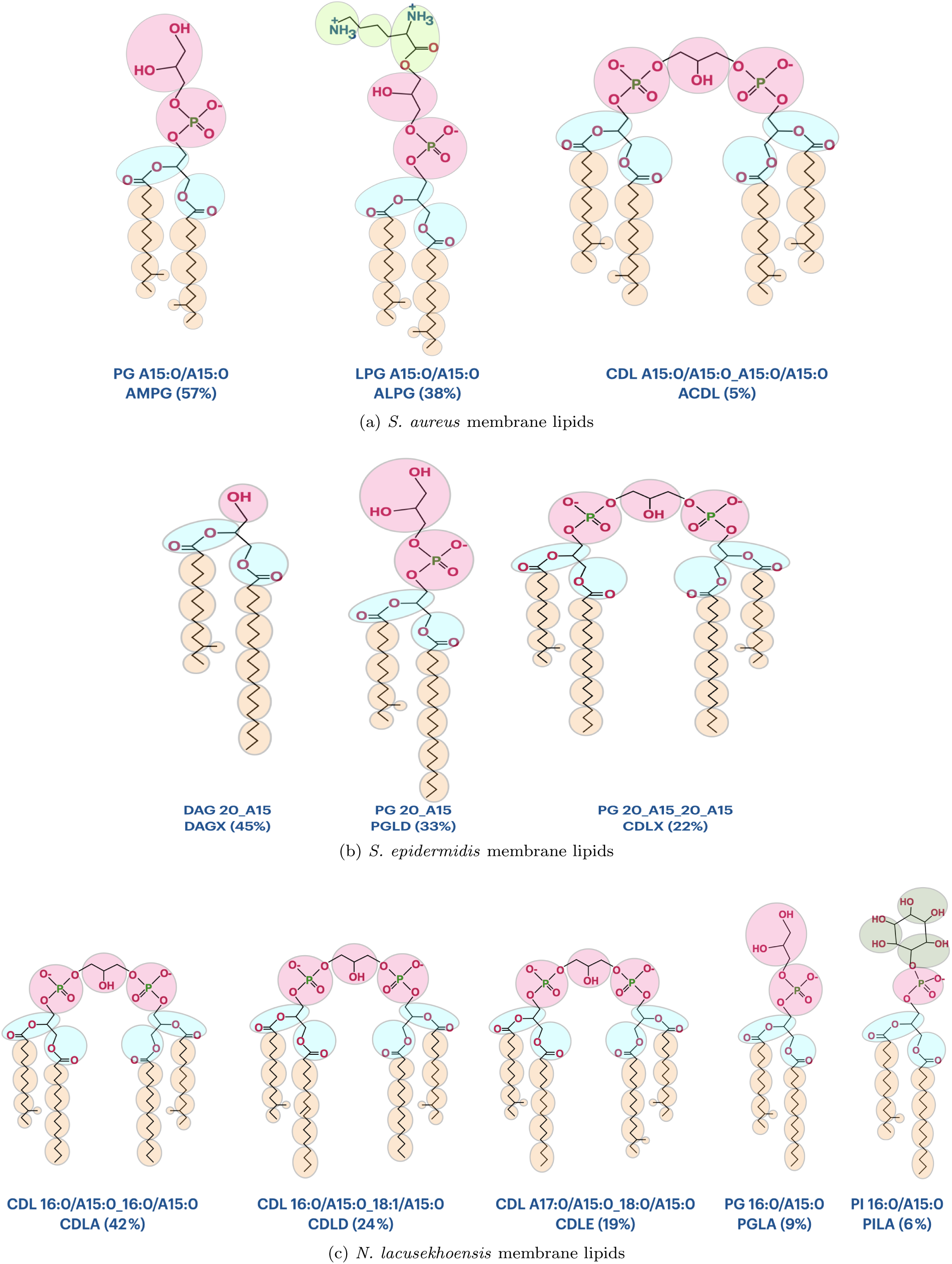
Lipid structures with percentage compositions for (a) *S. aureus* (b) *S. epidermidis*, and (c) *N. lacusekhoensis* membranes. The coarse-grained beads are also shown with mapping and underlying atoms. The carbon chains with label A in lipid nomenclature indicate the anteiso branching in the chains.

**Table 1:**
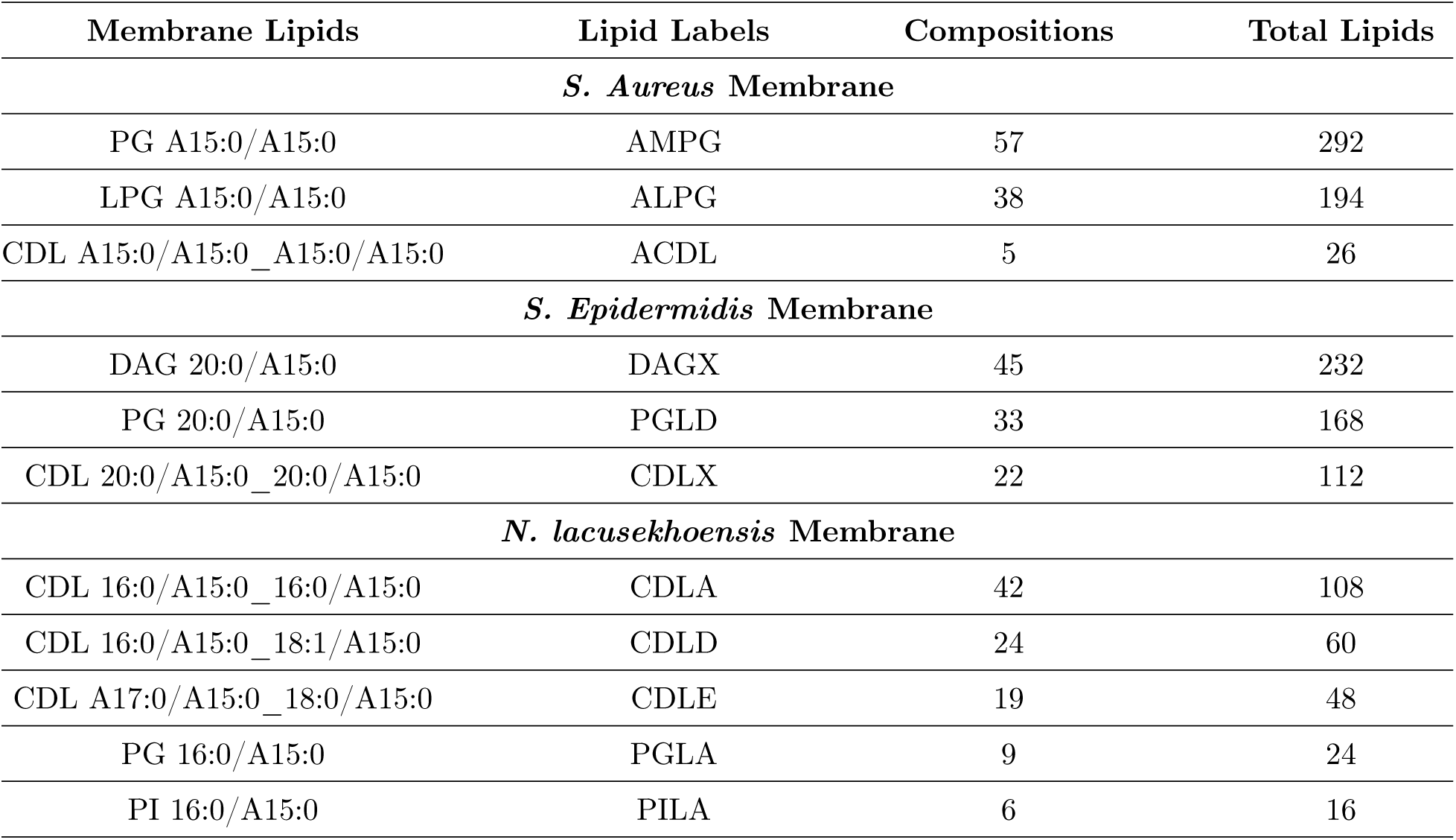
Membrane lipids and their molar compositions.

The force field parameters for lipids with atomistic details in our study are derived from the atom types and atomic charges of other phosphatidylglycerol lipids, cardiolipins and inositol lipid models available in CHARMM36 force field. ^29,30^ We first prepared SMILES structure codes and MOL2 files for these lipids, which were subsequently used to get lipid structures in pdb/gro format using CGenFF server. ^31^ The initial structures for lipid membranes were generated using MemGen server. ^32,33^ The coarse-grained structures for individual lipids as well as lipid membranes were created from corresponding atomistic systems, following the Martini-3 mapping schemes shown in Figure 2.

## 4 METHODOLOGY FOR MOLECULAR DYNAMICS SIMULATION

We have carried out all-atom (AA) and coarse-grained (CG) Martini simulations for lipids in bulk water and membranes using GROMACS suite 2022.5 at temperature 310 K and pressure 1 bar. The initial configurations for lipids and membranes were energy minimized using steepest descent algorithm in GROMACS, and the equations of motion were integrated using the leap-frog algorithm ^34^ in subsequent equilibration and production runs. For visualizing the simulation trajectories, we used the Visual Molecular Dynamics (VMD) software. ^35^

### 4.1 Atomistic Simulations

The systems with lipids in bulk water were equilibrated with a time step of 1 fs for 125 ps using Berendsen thermostat ^36^ followed by a 1 ns equilibration run using the Nosé-Hoover thermostat ^37^ with a time step of 2 fs. The time constant for the thermostat was 1 ps. The pressure was then maintained isotropically by employing the Berendsen barostat for 1 ns and Parrinello-Rahman ^38^ for another 500 ns acquisition run. The pressure coupling time constant of 5 ps and compressibility 4.5 × 10^−5^ bar^−1^ were used.

For membrane systems the lateral and normal pressure components were independently controlled using semiisotropic pressure coupling, and the production run was 1 *µ*s for each membrane. The hydrogen bonds were constrained using the LINCS algorithm. ^39^ The van der Waals forces were computed with a cutoff radius of 1.2 nm, and truncated gradually over distances in a range of 1-1.2 nm using a force-switch modifier. The Coulomb forces were computed using the Particle Mesh Ewald (PME) technique, with a cutoff radius of 1.2 nm for short range Coulomb interactions. ^40^

### 4.2 Coarse-grained Martini Simulation

The particle trajectories in CG simulations were computed at time step size of 20 fs. The temperature was maintained by rescaling the particle velocities using v-rescale thermostat ^41^ at coupling time constant of 1 ps. The barostat of Parrinello-Rahman was used for pressure control, with 12 ps time constant and 3 × 10^−4^ bar^−1^ compressibility. The nonbonded interactions were computed with a distance cutoff of 1.1 nm. The reaction-field method ^42,43^ was employed to solve long range Coulomb interactions using the relative dielectric constant 15. Each simulation trajectory is 2 *µ*s long for CG lipids in bulk water systems, and 10 *µ*s for CG membrane systems.

## 5 SIMULATIONS OF LIPIDS IN WATER

We have developed and parametrized the molecular models of membrane lipids for Gram-positive bacteria. The force field parameters for AA lipids are obtained from other phosphatidylglycerol and inositol lipid models in the CHARMM36 force field, and the parameters for the CG models were optimized by simulating the individual lipids in bulk water. The Lennard-Jones parameters for van der Waals interactions among the CG beads are defined by the bead types ascribed to the CG beads based on the underlying functional groups, following the standard guidelines of the Martini-3 force field. The bonded parameters, namely equilibrium bond lengths, angles and corresponding stiffness constants, are optimized by matching the CG distributions for bonds and angles with corresponding distributions obtained in AA simulations. The phosphate bead in phosphatidylinositol lipid is kept out-of-plane of the inositol ring using a dihedral potential. The procedure for optimizing bonded interaction parameters has been described in our earlier work. ^44^

Figure 3 compares the solvent accessible surface area (SASA) of the CG lipids with the corresponding AA structures. The comparison shows an excellent match of the SASA values between AA and CG models. The values are expected to be somewhat greater for quadruple-chain lipids when compared with lipids having two tails. The corresponding histograms for SASA are provided in Figure S2 of SI.

**Figure 3:**
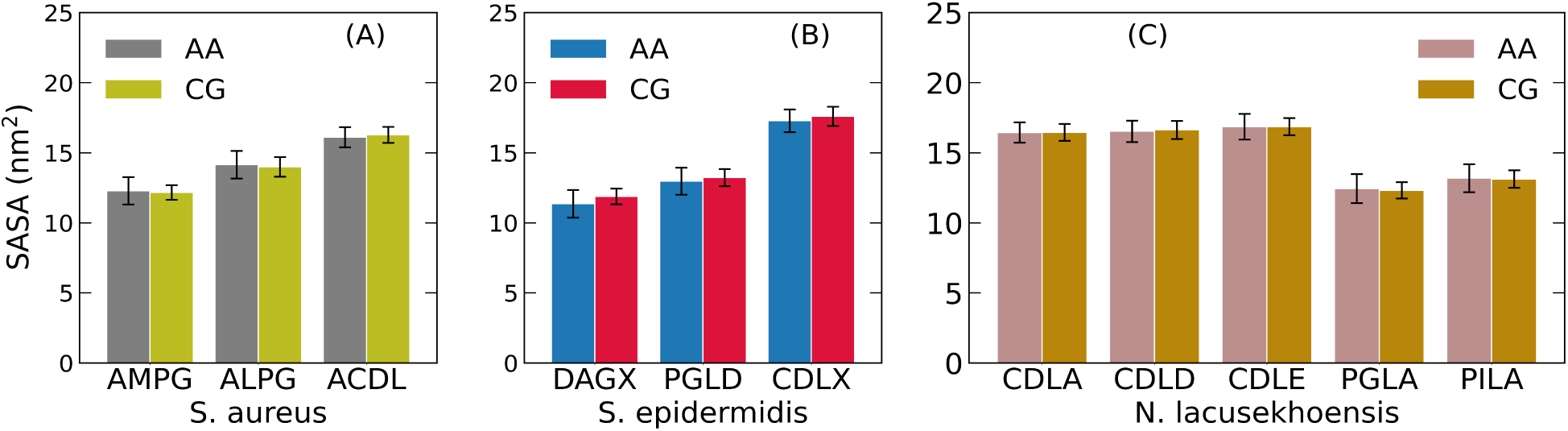
The solvent accessible surface area (SASA) for atomistic and coarse-grained models of membrane lipids in (A) *S. aureus*, (B) *S. epidermidis* and (C) *N. lacusekhoensis*. The different lipids for the three Gram-positive species are illustrated in Figure 2.

## 6 MEMBRANE SIMULATIONS

We have simulated cytoplasmic membranes of *S. aureus*, *N. lacusekhoensis* and *S. epidermidis* strains of Gram-positive bacteria. Lipid membranes with atomistic details were constructed using the MemGen server ^33^. The equivalent CG membranes were obtained by mapping the lipid atoms onto the CG martini beads using mapping schemes shown in Figure 2.

### 6.1 Membrane Area and Thickness

We have computed the structural properties of the bacterial membranes. The simulation box dimensions are used to evaluate the area per lipid tails, and the membrane thickness is computed based on distances between the phosphate headgroups along the membrane normal (Figure 4). Interestingly, despite the wide differences between the lipid headgroups and tail architecture the areas per lipid tails are quite similar between the different strains of Gram-positive bacteria. The lysyl lipids in *S. aureus* with bulkier lysine amino residue headgroups, effectively screens the apolar acyl chains from their interactions with polar water molecules. Consequently the carbon chains in *S. aureus* are relatively more flexible compared to the ordering of the acyl chains in the *S. epidermidis* membrane. However, the smaller headgroups of DAG lipids and a greater hydrophobic mismatch between carbon tails (20 and 15 carbon long chains) of *S. epidermidis* lipids facilitate a relatively tighter packing of lipids, resulting in a smaller area for the *S. epidermidis* membrane when compared with the other two strains. Longer carbon chains (20 carbons) in *S. epidermidis* result in a thicker membrane when compared with *S. aureus* and *N. lacusekhoensis* membranes, which are mostly populated with lipids of 15 and 16 carbon chains. The thicker membrane for *S. epidermidis* is also consistent with the lower area per lipid tail, which is an indication of improved lipid packing.

**Figure 4:**
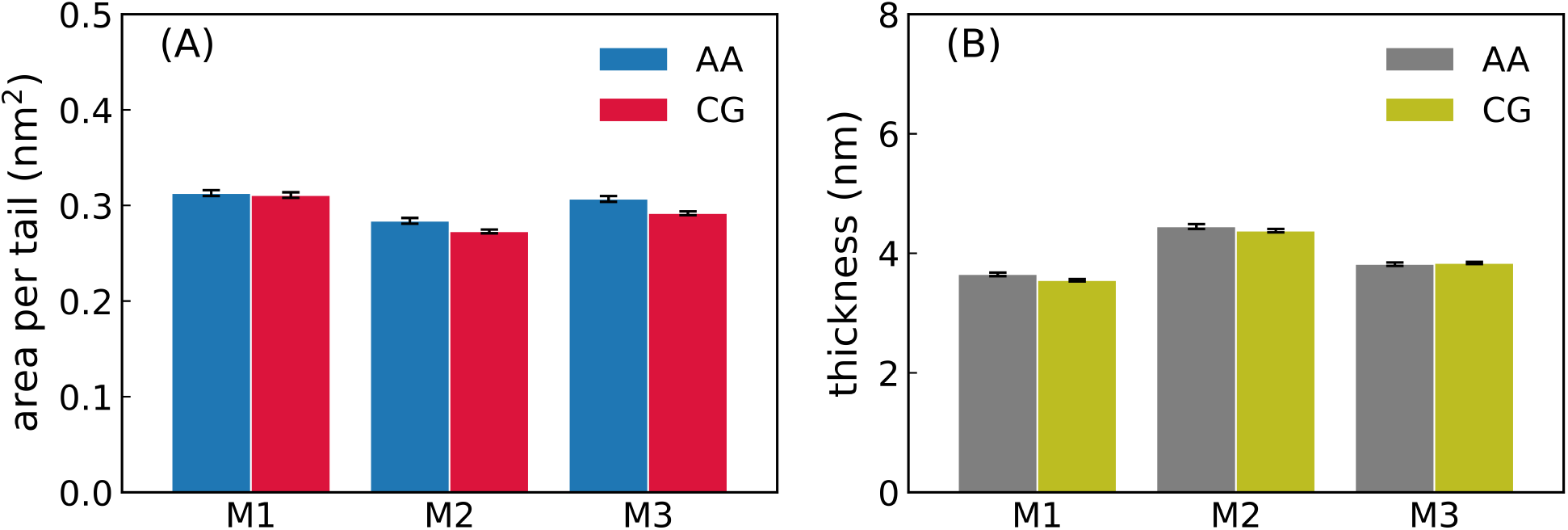
Comparison of (A) membrane area, and (B) membrane thickness for the three bacterial membranes, namely *S. aureus* (M1), *S. epidermidis* (M2) and *N. lacusekhoensis* (M3).

To quantify the acyl chain order parameters and phase of the lipid bilayers, we computed for atomistic membranes the deuterium order parameters, 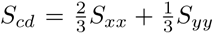. Here 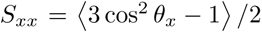, where *θ_x_* is the angle between the membrane normal (z-axis) and the x-axis within an appropriate molecular frame of reference. ^45^ The lipids in all three membranes follow similar trends in tail ordering along the carbon chains (Figure S3 in SI). A distinct increase in chain order is observed for *S. epidermidis* with the lowest ordering observed for *S. aureus* particularly near the headgroup region. These trends in acyl chain order are consistent with the variations in lipid tail area and membrane thickness variations between the strains (Figure 4).

### 6.2 Membrane Density and Ion Distribution

The comparison of density profiles between atomistic and CG simulation models is excellent for the model membranes of the three Gram-positive bacterial strains (Figure 5). The densities for water, ions, and lipids are accurately reproduced by the Martini-3 CG model membranes. On account of differences in masses of CG beads representing the lipid tails and the corresponding underlying atoms, there are differences in densities of CG lipids and the corresponding AA lipids in the core of the membrane (−2 < z < 2). The peaks for phosphate and lysyl groups in CG simulations also match with corresponding peaks obtained in atomistic simulations, indicating accurate positioning of the headgroups relative to the membrane center (Figure 5 D-F). Since the *S. epidermidis* membrane is dominated by DAG lipids, which are neutral, the concentration of potassium ions near the lipid headgroups is relatively low when compared with potassium ion concentration for *N. lacusekhoensis*, which shows the highest potassium density among the different strains. The increase in potassium ion density with cardiolipin content is clearly observed in the density profiles across the three strains and the sharper density peaks in *N. lacusekhoensis* indicates that the ions are tightly bound to the lipid headgroups. This ability to have a strongly bound counterion layer reflects a mechanism by which halophilic bacterial strains survive in highly saline environments by keeping the headgroups hydrated. ^46^ Furthermore, the conical shape of cardiolipin helps retain alkyl chain fluidity for the bacteria to survive, despite high counterion condensation. ^47^

**Figure 5:**
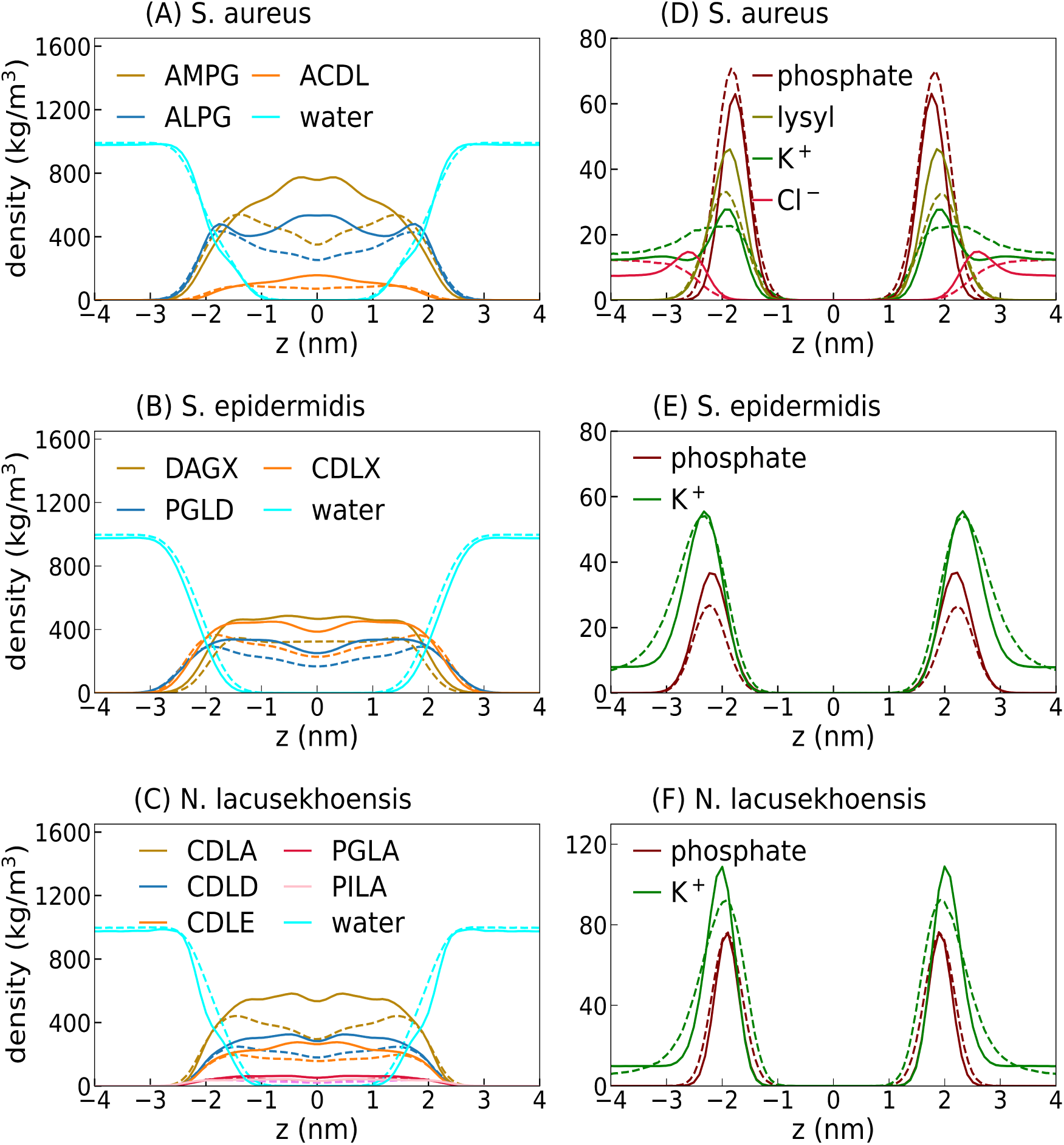
Comparison of density profiles for membrane lipids, ions and water in *S. aureus*, *S. epidermidis* and *N. lacusekhoensis*. The dashed lines are from the all-atom simulations and the solid lines correspond to the CG simulations. The phosphate and lysyl densities in panels D-F are scaled down by a factor 5 for clarity.

Figure 6 depicts the coordination numbers for ions around the phosphorus atoms in AA and phosphate beads in CG, confirming the accuracy and choice of the parametrized CG models. The hydration of lipid headgroups and distribution of ions around the charged groups are precisely captured, and this fact is further evident from the radial distribution functions for headgroup-ions (Figure 6), and headgroup-water (Figure S4 in SI). This ability of the Martini-3 force field to capture ion hydration and binding effects was also observed in our earlier parametrizaton of the Gram-negative lipopolysaccharide membrane. ^44^

**Figure 6:**
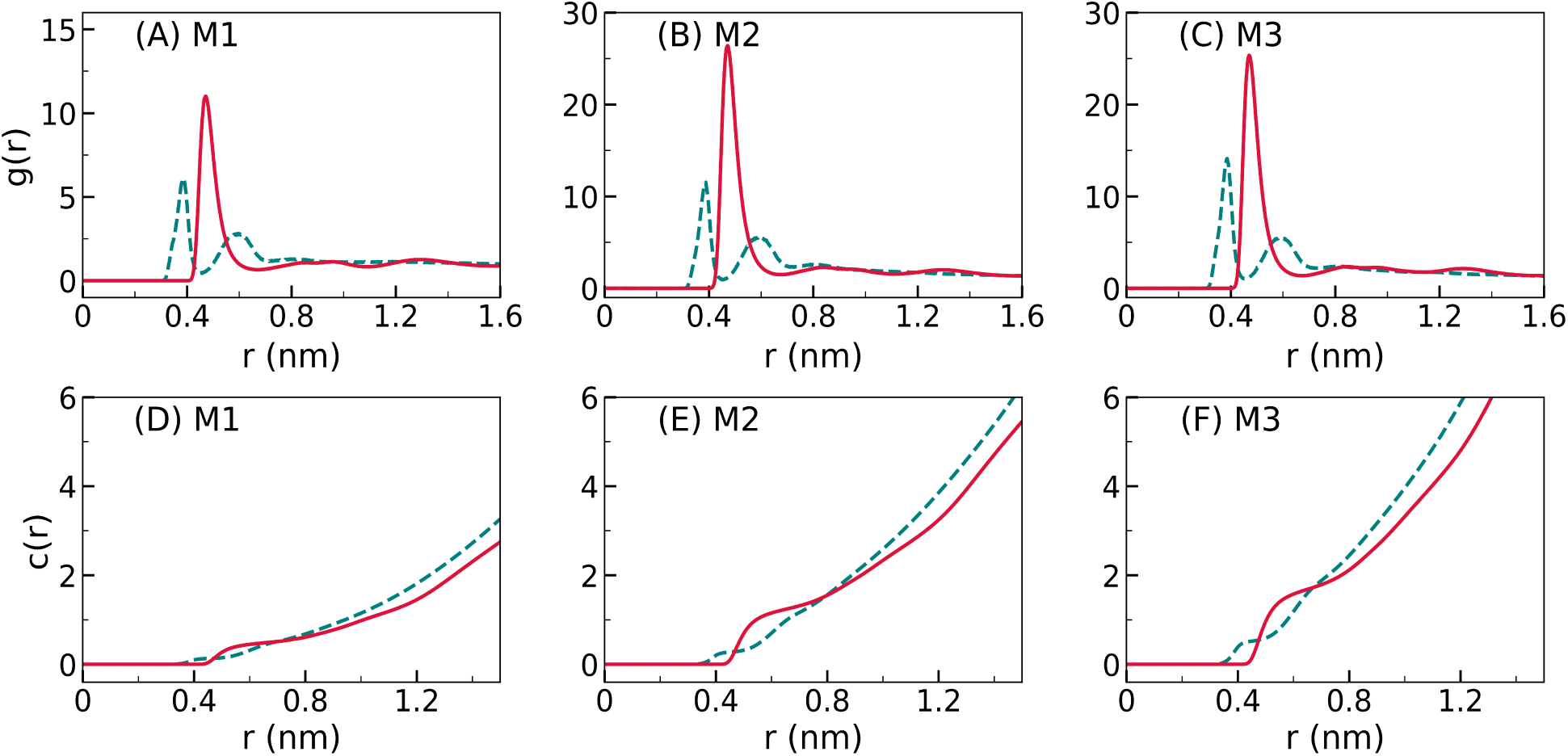
(A-C) Radial distribution functions indicating local structuring of potassium ions near the phosphorus atoms in AA and phosphate beads in CG membranes for *S. aureus* (M1), *S. epidermidis* (M2) and *N. lacusekhoensis* (M3) membranes. (D-F) The corresponding coordination numbers for potassium ions around the headgroups. The dotted lines refer to the AA simulations, while the solid lines correspond to the CG simulations.

### 6.3 Membrane Charge Distribution

The charge environment near the lipid-water interfaces plays a crucial role in lipid packing, headgroup hydration, membrane permeability and its resistance against antimicrobial molecules. Figure 7 shows the charge density (including counterions) obtained from CG membrane simulations. It is evident that the *N. lacusekhoensis* membrane is relatively more anionic when compared with the other two strains. The positive charge density at the lipid-water interfaces in the *S. aureus* membrane is attributed to the cationic charges of lysyl lipids in *S. aureus* (Figure 2A). In absence of cationic lipids in *N. lacusekhoensis* and *S. epidermidis*, the charge density profiles for these membranes result from the purely anionic nature of the lipids. Consistent with the densities of various charge moieties shown in Figure 5 (D-F), the overall charge densities at the lipid-water interfaces show peaks due to potassium ions (in addition to lysyl headgroups in *S. aureus*) surrounding the phosphate headgroups. A relatively high concentration of potassium for *N. lacusekhoensis* (Figure 5F) in comparison to *S. epidermidis* results in higher positive charge densities at the lipid-water interfaces in *N. lacusekhoensis*. These distinctive features will have a contrasting impact on interactions of the model membranes with charged molecules and peptides as we will illusrate later in the text.

**Figure 7:**
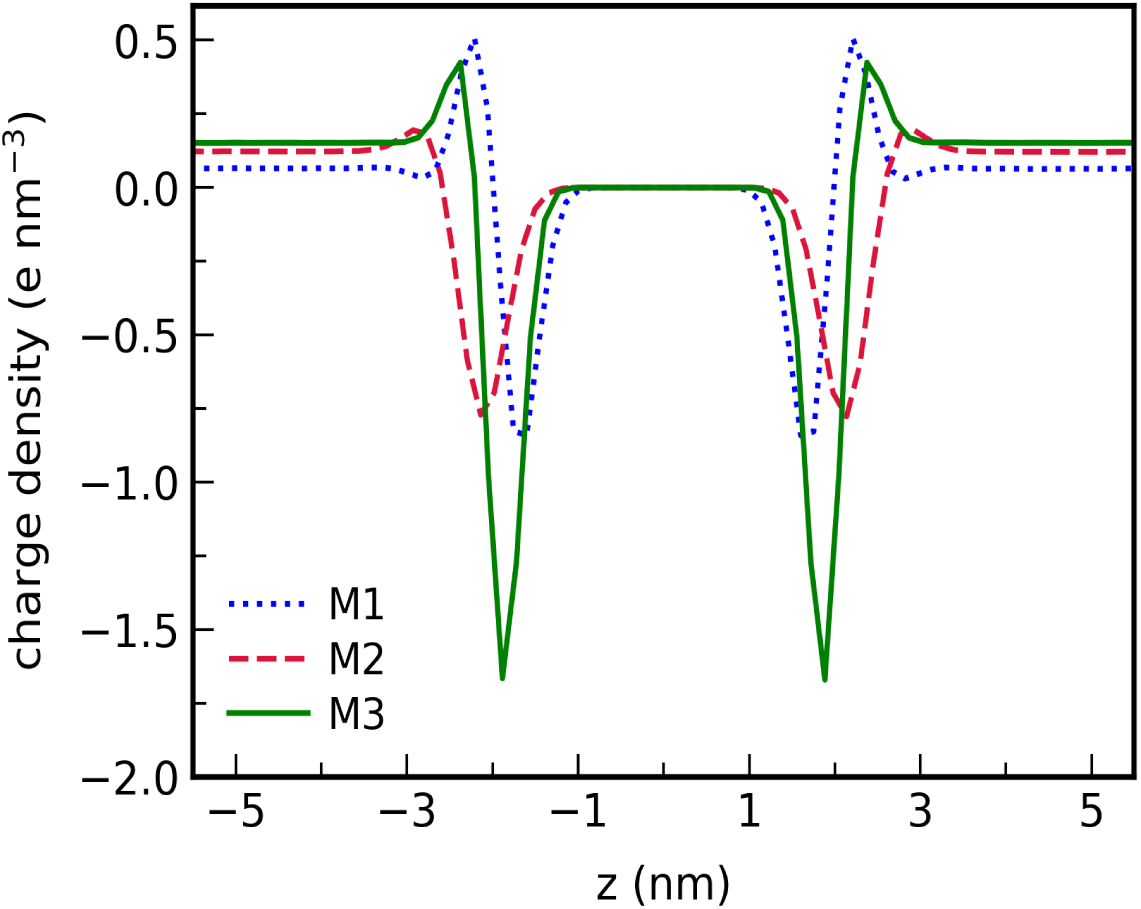
The charge density profiles for CG membrane systems arising from the charged moieties of lipids and ions across the membrane normal. The enhanced positive charge densities at the lipid-water interfaces are due to lysyl headgroups in *S. aureus* membrane and a high counterion density for *N. lacusekhoensis* (Figure 5D-F)

### 6.4 Lipid Mixing and Lateral Diffusion

In multicomponent membranes, the lipids having contrasting interactions can result in domain formation. In order to quantify the lipid mixing in the different Gram-positive strains, we have calculated the fractions of lipid-lipid interactions. The neighbor list for each lipid-lipid interaction is prepared based on the x-y coordinates of the center-of-mass positions of lipids in each leaflet, following the procedure by De et al. ^48^ The lipid-lipid fractional interaction parameter,

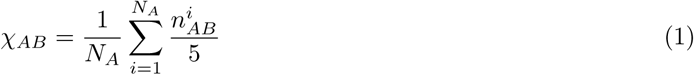

where *N_A_* is the number of lipids of type *A* and 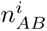 is the number of first five nearest neighbours of type *B* centered around *A*. Figure 8 depicts the fractional lipid-lipid interactions, which is akin to a lipid-lipid contact map, indicative of the relative affinity of a lipid to remain surrounded by another type of lipid. The reported fractions are averaged over the simulation trajectories. The affinity of AMPG with ALPG (0.61) in *S. aureus* is a bit stronger compared to AMPG-AMPG (0.55). Similarly DAGX lipid in *S. epidermidis* shows higher fractions with CDLX lipid (0.52) in comparison to DAGX and PGLD. While in *N. lacusekhoensis* the different cardiolipin molecules of varying length have similar trends of interactions. The decreasing trends in *χ_AB_* values in each column in Figure 8 are proportional to the lipid compositions in the CG membranes.

**Figure 8:**
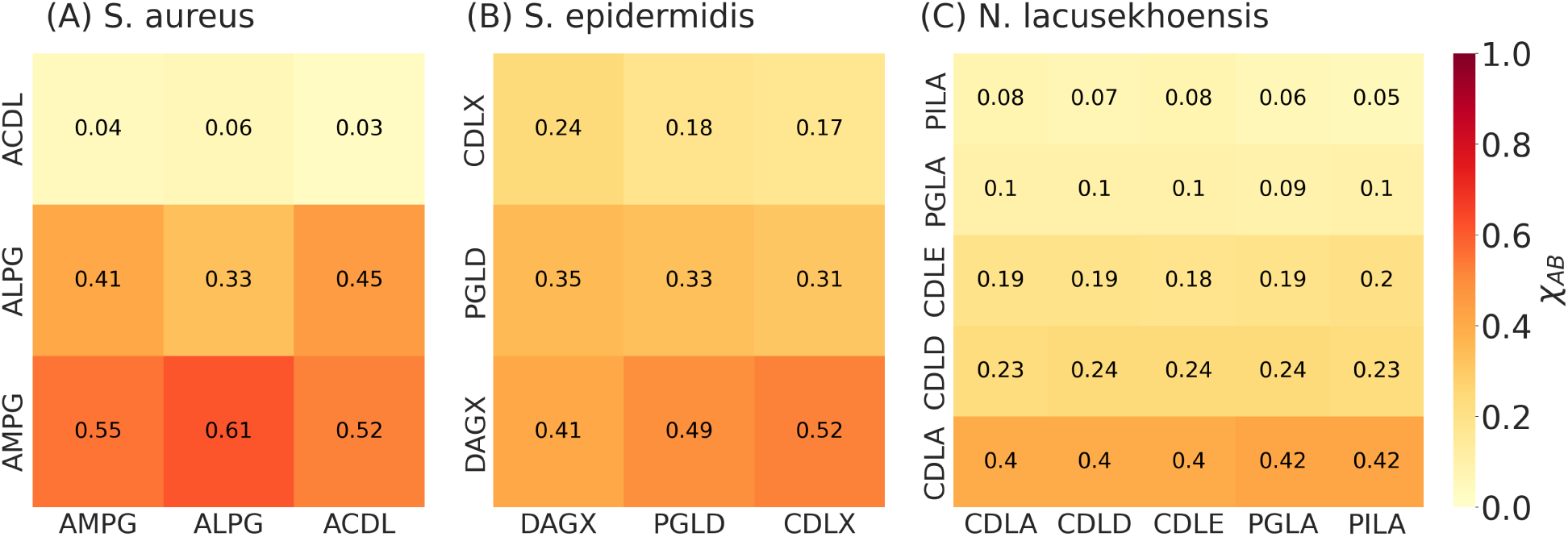
Number fraction *χ_AB_* of B type of lipid (y-label) found in the nearest neighbors of lipid type A (x-label) in CG membranes.

Figures S5-S7 given in SI show a comparison of the histograms for fractions of one type of lipid in nearest vicinity of another type of lipid in atomistic membrane simulations with those obtained in corresponding CG simulations. An excellent agreement between CG and AA data provides further validation of the choice for CG beads, which accurately capture the lipid-lipid nonbonded interactions. This illustrates that the lipophilic contacts among the lipids in atomistic simulations are precisely preserved by the CG lipid models developed for the Gram-positive membranes in our study.

The dynamics of lipid motion is assessed by computing the mean squared displacement (MSD) of lipids over a 10 *µ*s trajectory, in the plane of membrane, as depicted in Figure S8 in SI. There is no significant difference in the lateral diffusion coefficient among the CG membrane lipids modelled in this work. We observe a distinct sub-diffusive regime up to 10 ns for all the strains. The model cardiolipin molecules exhibited slightly slowest diffusion, while the inositol lipid (PILA) showed a little enhanced diffusion compared to other membrane lipids. We used the MSD data in time interval (t) 100-300 ns to extract the diffusion coefficient (D) by fitting in-plane diffusion to MSD = 4Dt. Nevertheless, the magnitude of diffusion coefficient ∼ 3 × 10^−7^ cm^2^*/*s is in agreement with other phospholipids reported in other simulations and experimental studies. ^49–51^

### 6.5 Mechanical Properties: Area Stretch and Bending Modulus

Mechanical properties of lipid membranes, such as area stretch modulus (*K_A_*) and bending modulus (*κ*), are important measures of the membrane energetics in response to structural changes during areal expansion and membrane bending. We computed the stretch modulus from the fluctuations of the lateral area using the simulation box dimensions. With increase in cardiolipin content a distinct increasing trend in the stretch modulus is observed (Figure 9A). *K_A_* varies from ∼ 370 mN/m in *S. aureus* to ∼ 475 mN/m in *N. lacusekhoensis*, reflecting a ∼ 28 % increase. The increasing trend observed for area stretch modulus with increasing cardiolipin content from 5% in *S. aureus* to 85% in *N. lacusekhoensis* supports previous observations on cardiolipin membranes. ^21^

**Figure 9:**
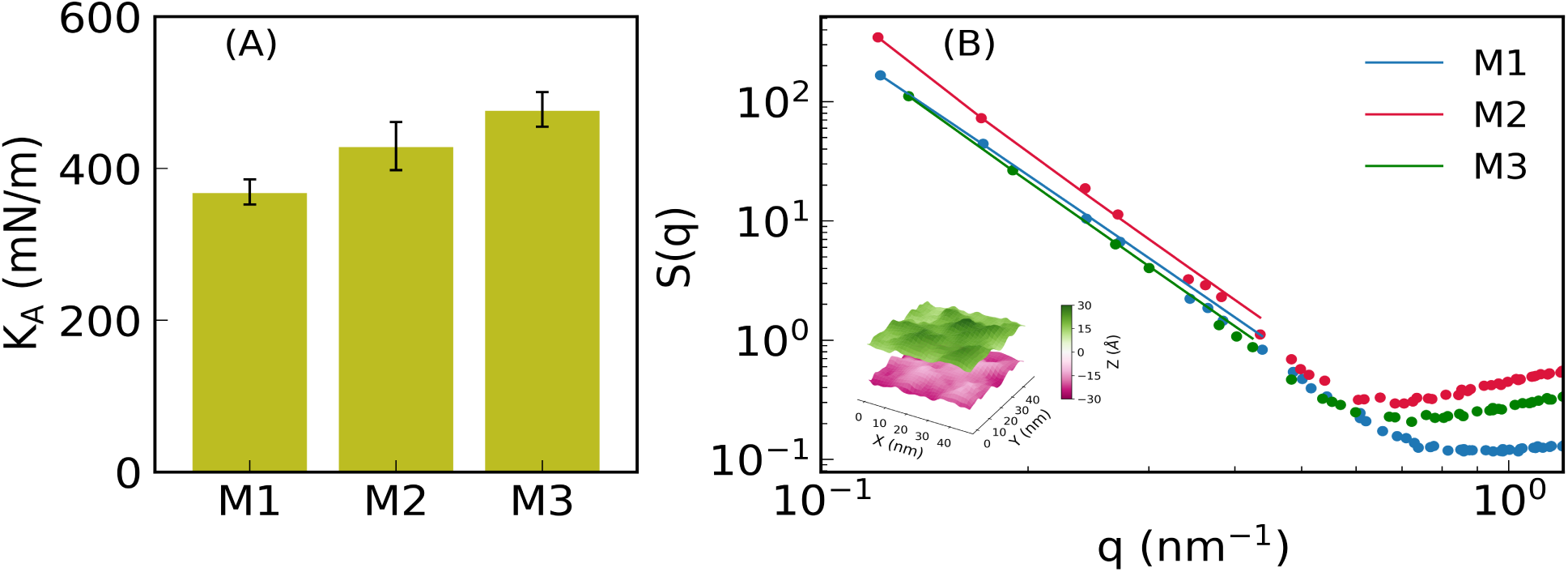
(A) The area stretch modulus for CG membranes. The trajectory averages and standard deviations are computed by splitting the simulation data into 50 blocks. The labels M1, M2 and M3 refer to *S. aureus*, *S. epidermidis* and *N. lacusekhoensis* membranes, respectively. (B) The membrane height fluctuation spectrum for CG membranes showing 1*/q*^4^ scaling at low wave vector q. The spatial variation of the membrane height along the membrane surface is indicated in the inset for a configuration taken from the *N. lacusekhoensis* membrane simulation.

To compute the bending modulus we constructed the CG Gram-positive membranes with lateral size ∼50 nm × 50 nm and simulated each membrane system for 3 *µ*s. These larger patches of membranes, which have 16 times more lipids than the lipids in relatively smaller membrane patches (Table 1), facilitate to capture the undulatory modes, especially at smaller wave vectors q=(2*πn_x_/L_x_*, 2*πn_y_/L_y_*). Here *n_x_* and *n_y_* are the integer numbers, and the lateral dimensions of membrane are denoted by *L_x_* and *L_y_*. According to Helfrich’s theory of membrane bending, ^52,53^ the structure factor, *S*(*q*) is related to the bending modulus *κ* and membrane tension *γ* as per *S*(*q*) = *k_B_T/*(*κq*^4^ + *γq*^2^). Here *k_B_* is the Boltzmann constant and *T* denotes temperature. We computed the structure factor *S*(*q*) using the fluctuations in membrane height,^54,55^ and the profiles for different Gram-positive membranes are depicted in Figure 9B. The structure factor data is fitted to *k_B_T/*(*κq*^4^ + *γq*^2^) in low *q* regime (*q* < 1 nm^−1^), and the bending modulus, *κ* is obtained from the fits. ^56,57^ The membrane tension values are vanishingly small (∼ 0.1 − 0.3 mN/m). The bending modulus values in our CG simulations are in a range of 19-30 *k_B_T*, with *S. epidermidis* membrane being more flexible with *κ* = 19.2 ± 0.7 *k_B_T*. The bending rigidity of *N. lacusekhoensis* (*κ* = 29 ± 0.7 *k_B_T*) and *S. aureus* (*κ* = 25±0.6 *k_B_T*) membranes are comparable. The cardiolipin molecule is diphosphatidyl glycerol, therefore its molecular architecture is similar to the phosphatidyl glycerol, with both having identical headgroups containing phosphate and glycerol. This gives rise to a comparable level of hydrophilic interactions of lipids with water at the membrane-water interfaces, which in turn govern the membrane undulations. Our results indicate that *N. lacusekhoensis* with the highest cardiolipin content has the highest *κ*. Interestingly despite its low cardiolipin content *S. aureus* membrane has bending modulus only marginally lower by ∼ 13%. It is known that cardiolipin concentration is enhanced in rod-like bacteria at the poles and septal regions cell membrane, where the local enhancement *κ* stabilizes the membrane deformation. ^58,59^ The varying charge density and different lipid types, which are known to influence mechanical properties, ^60–62^ complicate a direct comparison between the different strains with standard scaling of *κ* with the membrane thickness. ^63^ The *S. epidermidis* membrane is rich in DAG, which undergoes rapid flip-flop events across the bilayer leaflets. This could also be responsible for rendering the membrane more flexible for bending and softer for areal expansion as has been observed with cholesterol flip-flop events in other lipid membranes. ^64,65^

## 7 INTERACTIONS WITH ANTIMICROBIALS AND PRESERVATIVES

In order to study the interactions of the Gram-positive membranes with antimicrobials and preservative molecules, we computed the insertion free energies for representative molecules. These are thymol, methylparaben (MEPB), ethyl-lauroyl-arginate (ELAR) and cecropin-melittin-15 (CM15) for the CG membranes. The molecular structures and the CG Martini-3 mappings for these molecules are shown in Figure 10. CM15 was coarse-grained using the standard Martinize script and elastic dynamic network constraints were not imposed. Thymol is a naturally occurring antimicrobial molecule, and readily translocates through bacterial membranes. ^66,67^ MEPB is a non-ionic molecule, and it has been widely studied as a preservative against bacteria in foods and cosmetic formulations. ^68–71^ ELAR is a cationic antibacterial molecule formed by amidation of lauric acid with ethyl arginate. ^72^ In contrast, CM15 is an antimicrobial peptide, and there is a growing interest in understanding the conformational changes associated with CM15 and other antimicrobial peptides interacting with bacterial membranes. ^13,73,74^ We employed umbrella sampling simulations for computing free energies. Distances over a range of 0-4.4 nm between the center-of-mass of the test molecule and the lipid membrane along the membrane normal (z) were scanned with a window size of 0.1 nm. Each umbrella window is simulated for 400 ns, and the test molecule is restrained using the harmonic potential of strength 1000 kJ mol^−1^ nm^−2^. The free energy profiles were converged within 200 ns run time at each window. The other simulation parameters for thermostat, barostat and interactions are similar to those used in vanilla MD simulations. For CM15, which has a virtual martini bead, simulations were carried out at lower time step of 10 fs and pressure was maintained using the C-rescale barostat with a coupling time constant of 20 ps.

**Figure 10:**
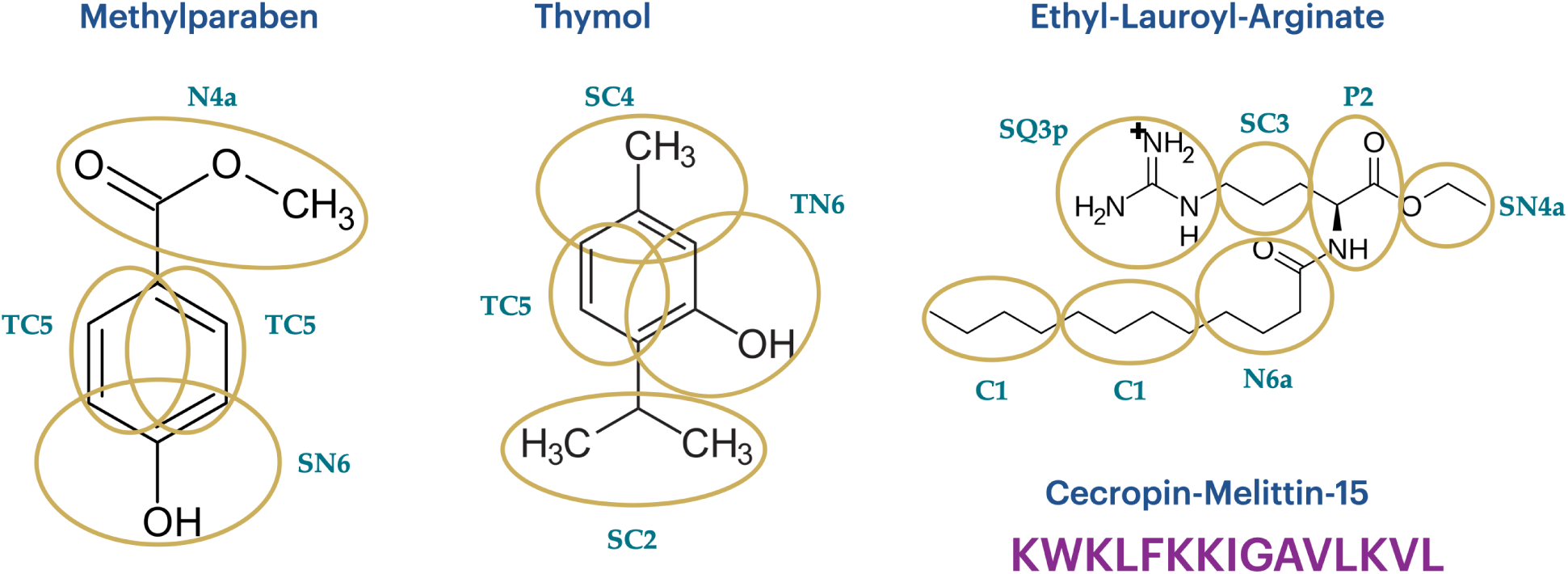
The atomic structures and coarse-grained mapping schemes for the methylparaben, thymol, and ethyl-lauroyl-arginate, as well as the peptide sequence for CM15. The bead types corresponding to Martini-3 force field are also indicated.

Figure 11 shows the potential of mean force (PMF) between the various molecules and the bacterial membranes. A general trend emerging from the PMF profiles is that the insertion free energy of electrostatically neutral molecules such as thymol and MEP are comparable across the three Gram-positive membranes. The small entry barriers (∼ 1 *k_B_T*) and their locations within the membrane for thymol (Figure 11A) and MEPB (Figure 11B) observed for *S. aureus* and *N. lacusekhoensis* membranes correlate with the enhanced positive charge densities (Figure 7) near the headgroups of these membranes. In contrast, a relatively uniform surface charge density for *S. epidermidis* did not result in a barrier at the membrane interface. The translocation free energy (energy required to cross the membrane center) of both thymol and MEPB are highest for the *S. aureus* membrane (Figures 11 A and B) when compared with the other two bacerial strains. This is consistent with higher lipid density at the center of *S. aureus* membrane due to lipids having symmetric chain lengths of 15 carbons in *S. aureus* (Figure 5A). In contrast, the hydrocarbon chains for all phospholipids components in both *S. epidermidis* and *N. lacusekhoensis* are asymmetric with chain lengths ranging from 15 to 20 carbons (Figure 2) resulting in a lower lipid density (Figures 5B and C) at the membrane interior allowing small molecules to translocate with lower energy barriers (Figures 11A and B). We note however, that this difference in translocation free energy is within a few *k_B_T*. Furthermore, MEPB with a higher extent of polar groups has a much greater translocation free energy when compared with thymol.

**Figure 11:**
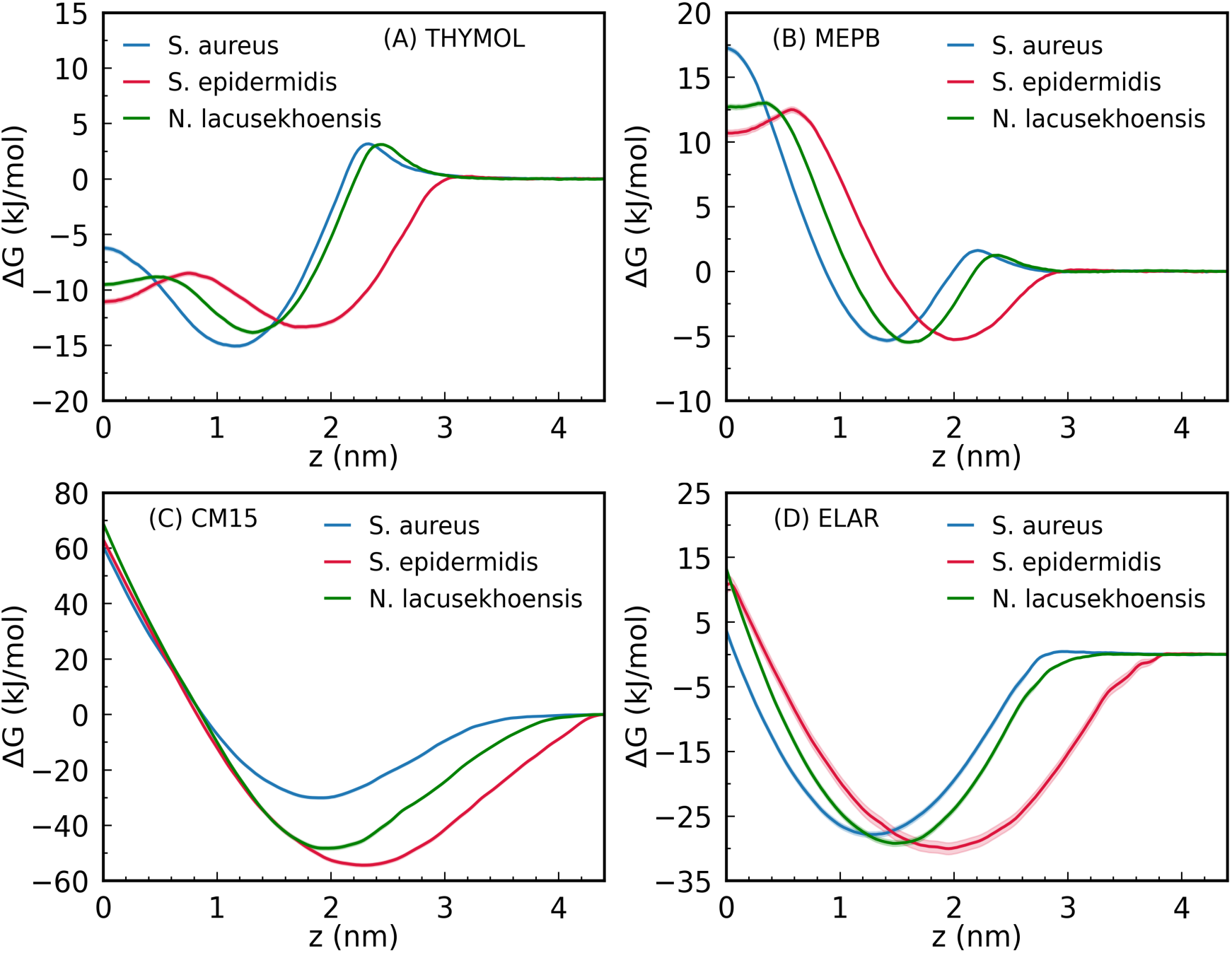
Potential of mean force between the bacterial membranes and test molecules, (A) thymol (B) methylparaben (C) cecropin-melittin-15 (D) ethyl-lauroyl-arginate.

In contrast to the small molecule free energy profiles and their differences between the three strains, the cationic molecules, CM15 (+5) (Figure 11C) and ELAR (+1) (Figure 11D) have a more favourable free energy of membrane partitioning into the different strains. The translocation free energy barriers to cross the bilayer midplane are significantly higher for these molecules when compared with thymol. CM15 is found to have a favourable free energy minima (8-20 *k_B_T*) for all the three strains, and the minima are located in the vicinity of the phospholipid headgroups. Here again, the local charge density at the headgroup interface (Figure 7) modulates the location and the magnitude of minima in free energy profiles. The most favourable insertion free energy is observed for *S. epidermidis*, which has the lowest positive charge at the headgroup region and the highest free energy occurs for *S. aureus*, where the highest positive charge density is observed. Since CM15 is known to undergo conformational changes upon binding and insertion, we present a more detailed analysis for CM15 along the free energy path. ^13,73,75,76^

We track the conformational change in CM15 by computing the end-to-end distance (l*_ee_*) between the terminal CG backbone beads LYS-1 and LEU-15 (Figure 12A) and the peptide’s orientation angle (*θ*) made by the vector connecting the terminal backbone beads with the membrane outward normal (z-axis). The distance l*_ee_* remains around a value of 2 nm for z *>* 2 nm and decreases to about 1.5 nm as the peptide approaches the membrane center in the repulsive region of the free energy. The angle *θ* shown in Figure 12B undergoes a large change as the peptide enters the headgroup region of the membrane, indicative of significant change in peptide orientation upon membrane entry. We observe a distinct rise in the angle from about 30° to 90° as the peptide starts interacting with membrane interface, implying a membrane perpendicular entry pathway with a stretched out peptide (high l*_ee_*) configuration (Figure 12C). However upon entry, the peptide prefers to adopt a membrane parallel orientation below the phospholipid headgroups. Interestingly, a similar entry mechanism is observed for all three strains despite the differing lipid compositions and charge content (Figures S9 and S10 in SI). In the repulsive part of the free energy, *θ* changes between 90-120 with a pronounced decrease in l*_ee_* due to a conformational change to a more collapsed state (Figure 12C). This region of the free energy is also correlated with large membrane undulations. CM15 is known to undergo a folding transition upon interacting with phospholipid membranes with a propensity to adopt a membrane parallel configuration below the phospholipid headgroups. Although the CG model cannot directly capture a folding event, our results are consistent with this mechanism. ^73,75,76^

**Figure 12:**
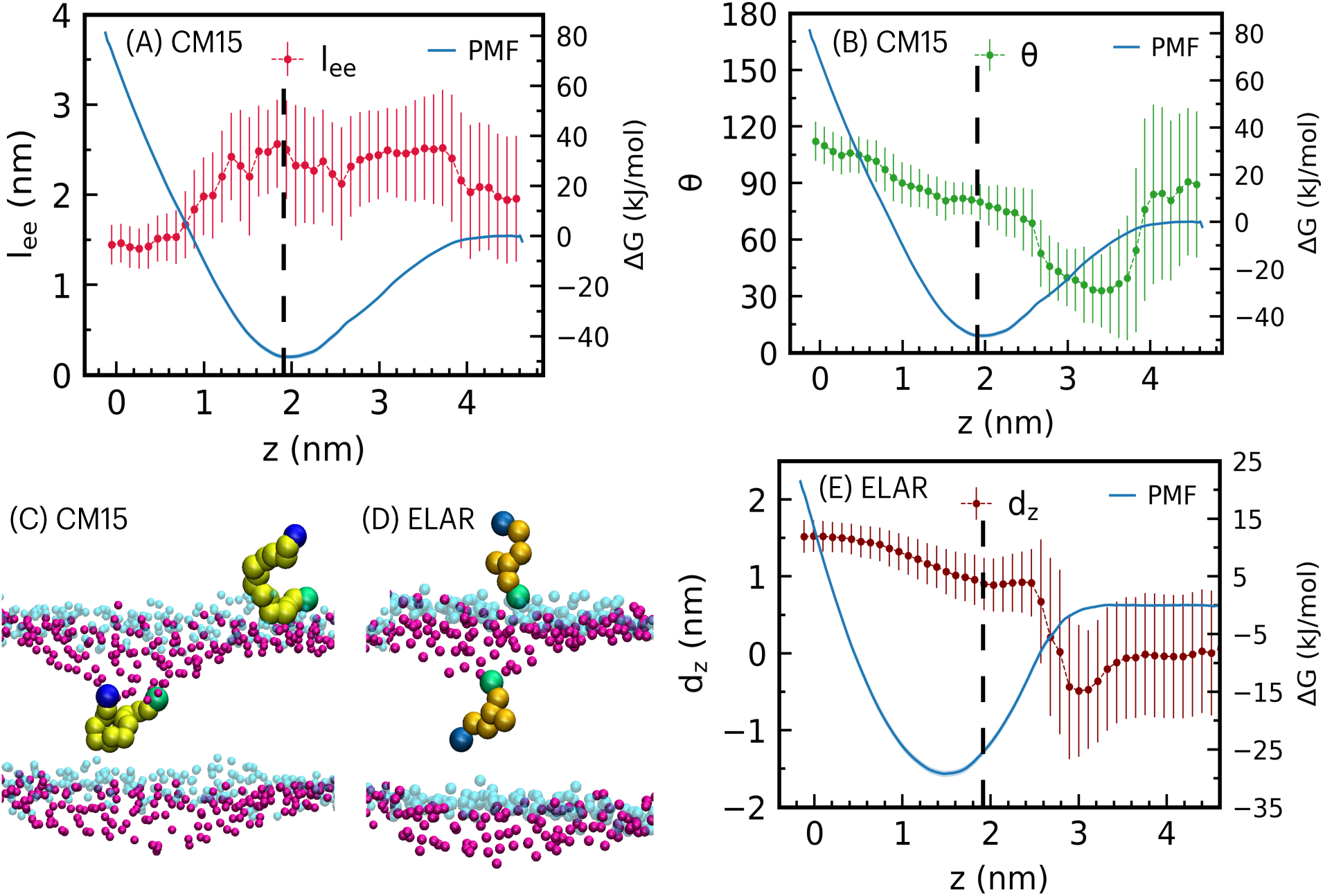
(A-C) Conformational changes of CM15 peptide interacting with *N. lacusekhoensis* CG membrane. (A) Peptide end-to-end distance (l*_ee_*) as peptide approaches the membrane center. (B) The angle (*θ*) variation with distance z. The angle is defined between the vector connecting terminal backbone beads and the membrane normal (z-axis). (C-D) The simulation snapshots were taken at instances when the molecule was outside the membrane (headgroups in translucent phosphate beads shown in cyan color) and when it was at membrane center (phosphate headgroup beads in crimson color). The peptide backbone beads are indicated by lime color with N- and C-terminal beads shown in green and blue colors, respectively. The charged amine group and terminal carbon bead of the acyl chain of ELAR (Figure 10) are shown in green and blue colors, respectively. (E) The preferential orientation of ELAR interacting with CG membrane of *N. lacusekhoensis*. The y-axis label d*_z_* is the z-component of the distance between the charged amine bead and the terminal carbon bead of ELAR. In above panels, the free energy (Δ*G*) is also indicated as a function of molecule’s distance from the membrane center, and the error bars are the standard deviations of the data. The vertical dashed lines in panels A, B and E mark the location of membrane-water interface.

During its interactions with Gram-positive membranes, ELAR adopts a preferential orientation while inserting into the membrane. Here again we observed similar trends in the free energy differences as observed for CM15 and a repulsive free energy to translocate across the membrane midplane. The free energy minima ranges between 8-10 *k_B_T* for ELAR. To quantify the orientational change taking place across the membrane interface, we computed the distance between the SQ3p bead and the terminal carbon C1 bead of the lauroyl chain (Figure 10). Figure 12E shows the z-component of this distance (d*_z_*) which clearly depicts a stark variation across the membrane interface (2 < z < 3). The ELAR molecule in the extracellular space is isotropically solvated with no preferential orientation. However, upon interacting with membrane lipids, the positive amine group of ELAR prefers to be surrounded by negatively charged lipid headgroups, with the lauroyl chain away from the hydrophilic lipid headgroups. This preferential orientation is more prominent in *N. lacusekhoensis* on account of the higher negative charge density in comparison to the other two membranes (Figures S9 and S10 in SI). The membrane of *S. epidermidis* is mainly composed of neutral DAG lipids and hence it has weaker electrostatic interactions with the ELAR molecule at the membrane interface, as implied by the d_z_ values at the interface (Figure S10E). The *S. aureus* membrane shows moderate electrostatics with the ELAR headgroup at the membrane interface due to positively charged lysyl lipids (Figure S9E). Once the molecule is inside the membrane, the acyl chain of ELAR becomes more ordered, with the amine headgroup reoriented toward the lipid headgroups, as indicated in Figure 12D.

## 8 INTERACTIONS AT HIGH SALT AND LOW pH

In this section, we explore the halophilic and alkalophilic nature of *N. lacusekhoensis* and study how salt concentrations and protonation states of the lipids modulate interactions with thymol. The motivation is to understand how external conditions alter microbial molecule entry into the membranes. To capture the differences in protonation and associated counterion binding states, we carried out all-atom simulations on charged and neutral membranes of *N. lacusekhoensis* with and without added salt. In order to simulate an environment with acidic pH, one oxygen atom within each phosphate headgroup for all lipids are protonated (Figure 2). The corresponding partial charges for the neighboring atoms and their atom types are altered within the framework of the CHARMM36 force field. We refer to this as the neutral membrane. The native membrane with the original charge distribution is referred to as the charged membrane in this section. Initially, 100 molecules of thymol were equally distributed in the aqueous regions both above and below the membrane, and membrane partitioning was studied for the charged *N. lacusekhoensis* and neutral membranes with 0 and 0.8 M KCl concentrations. Regardless of the charge and salt concentration, all the thymol molecules eventually partitioned into the membrane, positioning themselves below the headgroups, as illustrated in Figure 13. Thymol orients with its weakly polar OH group facing toward the phospholipid headgroups and similar positioning for thymol was observed in Gram-negative *E. coli* membranes. ^77^ Due to the weakly polar nature of the thymol, the thymol density distribution is narrower in the charged membrane when compared with the neutral membrane (Figures 13A and B). The K^+^ counterion density peaks are observed only for the charged membrane and this plays a key role in modulating the kinetics of membrane partitioning.

**Figure 13:**
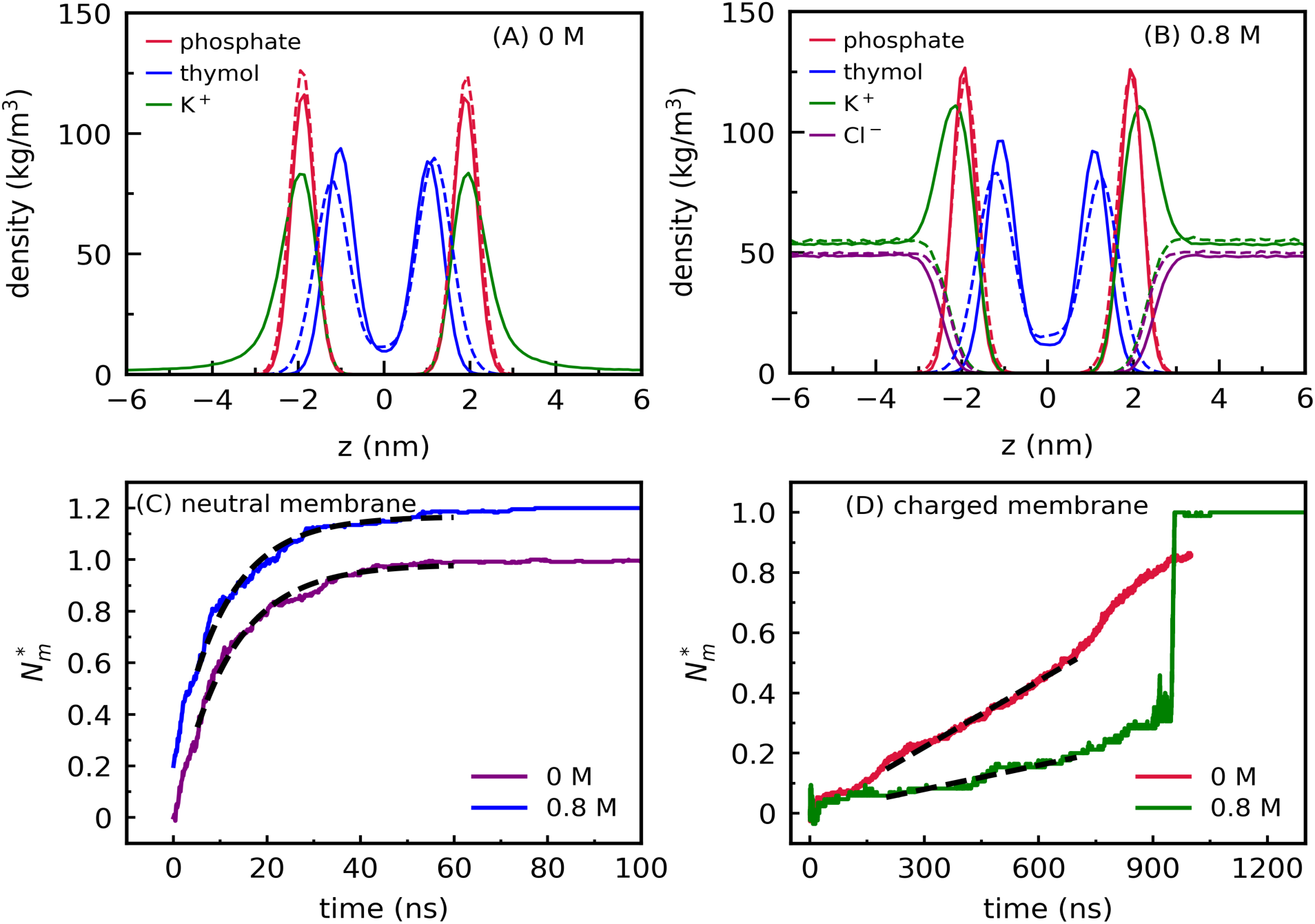
The density profiles for phosphate headgroups, thymol molecules and ions in all-atom membrane systems at (A) 0 M and (B) 0.8 M salt concentrations. The phosphate densities are scaled down by factor of 3 for visualization purpose. The dotted lines correspond to the neutral membranes and the solid lines indicate the profiles for charged membranes. (C-D) Fractions (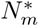) of thymol molecules in the neutral and charged AA membranes. The black dashed lines are the fits for (C) first and (D) zeroth order kinetics. The data for neutral membrane with salt (0.8 M) in panel (C) are shifted in y-axis by 0.2 for better visualization purpose.

We monitor the kinetics of membrane partitioning by tracking the number of thymol molecules that enter the membrane as a function of time. We define 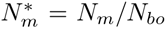, where *N_m_* is the number of thymol molecules in the membrane and *N_bo_* is the initial number of molecules in the bulk solution ^71^. In Figures 13C and D the time variation of 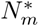 depicts distinct differences between the kinetics for the neutral and charged membranes. For the neutral membrane where counterion condensation at the membrane headgroups is not observed, a complete membrane partitioning is observed within ∼ 100 ns (Figures 13C and S11A). Over this time period no significant aggregates of thymol were formed in solution (Figure S12 in SI).

In contrast the partitioning time scale for the charged membrane is considerably slower, and complete partitioning takes ∼ 1*µ*s (Figures 13D and S11B). For the charged membrane, the cloud of K^+^ ions near the headgroups (Figures 13A and B) poses a barrier for thymol molecules to enter the membranes (Figure 11A). This entry barrier retards the kinetics of thymol partitioning into the charged membrane giving sufficient time for thymol molecules to form an aggregate in bulk solution. Aggregation is complete within ∼ 100 ns time (*t*), and subsequent partitioning of thymol occurs due to release of thymol molecules from the aggregate. Distinct jumps in 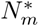 occur due to fusion of the aggregate with the membrane as observed in the 0.8 M case (Figure S11B). Similar fusion events of micelles were observed for small molecule preservatives, surfactants and ionic liquids with bacterial and phospholipid membranes. ^71,78^ This partitioning phenomenon can be observed in simulation snapshots shown for the charged membrane in Figure S13 of SI.

To quantify the different kinetic mechanisms we use a recently developed first order kinetic model for the neutral membrane and a zeroth order kinetic model for the charged membrane, respectively. ^71^ In the study by Chockalingam et al., ^71,78^ first order kinetics occurred when molecules in the bulk solution did not form aggregates, however zeroth order kinetics occurred when aggregates were present in solution. Slow sequential release of molecules from the aggregate gave rise to the zeroth order kinetics. We observe a similar mechanism governing the kinetics in this study, however the formation of thymol aggregates, as well as thymol release, is modulated by the presence of charge. We fit the data to 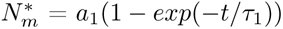 for the neutral membrane and obtain the first order time constant *τ*_1_ = ∼ 12 and ∼ 11 ns for the 0 M and 0.8 M salt conditions respectively. For the case of the charged (native) membrane we obtain the zeroth order time constant *τ_o_*= ∼ 1.4 *µs* and ∼ 3.8 *µs* using 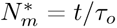, respectively for the 0 M and 0.8 M cases. These represent fits are averaged over three independent simulations, except for the charged membrane with salt case where we fit the individual simulation runs due to aggregate fusion with the membrane (Figures 13C and D).

The barrier and kinetics are governed by the concentration of counterions at the phosphate headgroups, their density correlations and the breakup kinetics of thymol from the aggregate. For the charged membrane, where thymol release from the aggregate controls the kinetics, our simulations suggest that the aggregates have greater stability at high salt concentration when compared to 0 M salt case. Both charged systems display zeroth order kinetics, however the rate of partitioning is slower at 0.8 M than 0 M case (Figures 13D and S11B) and we attribute this to counterion condensation at 0.8 M which leads to lowered headgroup-headgroup repulsion and enhanced lipid packing (Figure S14). Hence the salt has a more complex influence on the zeroth order kinetics. In addition to the enhanced counterion density and correlations with headgroups (Figure S11C) at the membrane interface, the kinetics of aggregate breakup and thymol release could also be influenced by the presence of salt.

In order to further validate these findings drawn from the atomistic membrane simulations, we computed insertion free energies for thymol at 0 and 0.8 M salt concentrations with charged and neutral CG membranes of *N. lacusekhoensis*. The free energy profiles are shown in Figure S11D in SI. The reference energy is set to zero in the aqueous regions away from the membrane. The negative charge on the phosphate headgroups in the charged CG membrane induces a net positive charge density of counterions near the headgroups (Figure 5F). As a result the positive charge density at the headgroups induces a small repulsive barrier in free energy, followed by a steep gradient for the molecule to enter into the membrane. In contrast, the free energy profiles for the neutral membrane with and without salt are relatively flat with no entry barrier at the phosphate headgroups due to absence of counterion density peaks (Figures 13A and B) near the headgroups. These trends are consistent with variations in kinetics observed earlier.

Before concluding this section, we briefly comment on the halophilic-alkophilic nature of *N. lacusekhoensis* which is known to survives in high salt and high pH environments. ^79,80^ Survival in high salt environments requires balancing internal osmotic pressure with osmolytes synthesized within cell or drawn from the extra-cellular milieu. ^46^ Additionally the membrane should retain its selective permeability under halophilic stress. Our simulations with thymol illustrate the retarded kinetics for the native (charged) membrane and lowered kinetics at high salt conditions (Figure 13D), akin to protective permeabilization as a result of counterion condensation on the cardiolipin headgroups. The results with the neutral membranes, mimics a situation of low pH or acidic environment where rapid kinetics of thymol uptake occurs as a result of lowered free energy barriers (Figure 13C).

## 9 SUMMARY AND CONCLUSIONS

We have carried out a detailed analysis of the inner membrane of three Gram-positive bacterial strains, namely *S. aureus*, *S. epidermidis* and *N. lacusekhoensis*. A lipidomics study of *S. epidermidis* and *N. lacusekhoensis* is used to obtain the lipid architecture and composition for these strains, which to our knowledge have not been reported in the published literature. Distinct differences between the strains arise from the lipid composition, charge and the cardiolipin variation from 5% in *S.aureus* to 85% in *N. lacusekhoensis*.

In order to study structure and mechanical properties of these widely different strains we have developed both atomistic and coarse-grained lipid membrane models. The coarse-grained Martini-3 model membranes are validated by reproducing the structural properties such as membrane area, thickness and density profiles with all-atom models. The ion correlations with the lipid headgroups are also in good agreement with the atomistic model membranes. The inter-lipid interactions dictate a homogeneous distribution of lipids in the membranes, indicating good lipid mixing in these widely varying multicomponent membranes. Lipid diffusion coefficients are comparable with other phospholipid membranes, and a distinct sub-diffusive regime is observed for all strains. The area stretch modulus increases with an increase in cardiolipin content, however although the highest bending modulus was observed for *N. lacusekhoensis*, a direct correlation with cardiolipin content was not observed.

The quantitative differences in the partitioning free energy profiles arise from the membrane charge, acyl chain length and charge distributions on the small molecule. Counterion condensation, together with the ion correlations, alters the charge density to modulate free energy barriers and kinetics of membrane partitioning. Using the CG membrane models we elucidate the entry mechanisms and free energy barriers for the antimicrobial peptide CM15, and a preservative molecule ELAR. Entry of CM15 is found to have a striking similarity across all three membranes with the molecule reorienting along the membrane normal prior to entry and adopting a membrane parallel configuration below the headgroups upon entry. Translocation free energies were the lowest for *S. aureus* and the greatest for *S. epidermidis*. In contrast, the translocation free energies for ELAR were quite similar across the three strains. Despite the tighter packing of lipids in *S. epidermidis*, its membrane is more fragile for insertion of the cationic molecules (ELR) and peptides (CM15), possibly due to DAG molecules, which undergo flip-flop events across the membrane leaflets.

We also illustrate the manner in which the halophilic and alkophilic strain, *N. lacusekhoensis* with its high cardiolipin content, modulates the partitioning kinetics of the antimcrobial molecule thymol with pH and salt.

In conclusion, the charge density modulations influences the free energy of insertion and specific differences in partitioning arise from the molecule charge and hydrophobic content. Interestingly, the membrane structural properties, bending rigidity and interactions with small molecules are largely conserved across the three model membranes, reflecting an ease of adaptability and survival of the bacterial species in harsh conditions of salt and pH despite the structural and compositional diversity in lipid environment for the different species.

## ACKNOWLEDGMENTS

We acknowledge Unilever Research and Development for funding and postdoctoral support. We also acknowledge Supercomputer Education and Research Center (SERC) at Indian Institute of Science for providing computational facilities under the National Supercomputing Mission.

## ASSOCIATED CONTENT

### Supporting Information

The Supporting Information is available. The table for lipidomics data; histograms for SASA; lipid chain order parameters; radial distribution functions (RDFs) and coordination numbers for water surrounding phosphorus atom; histograms for fractional lipid-lipid interactions; mean squared displacement data; peptide end-to-end distance and molecule orientation for interactions of CM15 and ELAR with *S. aureus* and *S. epidermidis* CG membranes; kinetics of thymol insertion into atomistic neutral as well as charged membranes of *N. lacusekhoensis* and corresponding coordination numbers for potassium ions around the phosphorus atom; thymol PMFs in charged and neutral membranes; simulation snapshots showing thymol insertion into neutral and charged membranes; lipid chain order parameters for charged *N. lacusekhoensis* membrane at 0 and 0.8 M salt concentrations.

## AUTHOR INFORMATION

### Author Contributions

R.V.: Setting up simulations, analysing trajectories, visualization, investigation, and manuscript writing and editing; E.C.: lipidomics data and manuscript writing and editing; M.W.: conceptualization, lipidomics data and manuscript writing and editing; K.G.A.: conceptualization, discussion, visualization, resources, manuscript writing and editing.

## References

[1] Christian Sohlenkamp and Otto Geiger. Bacterial membrane lipids: diversity in structures and pathways. FEMS Microbio. Rev., 40(1):133–159, 2016.

[2] John TJ Cheng, John D Hale, Melissa Elliott, Robert EW Hancock, and Suzana K Straus. The importance of bacterial membrane composition in the structure and function of aurein 2.2 and selected variants. Biochimica et Biophysica Acta (BBA)-Biomembranes, 1808(3):622–633, 2011.

[3] Pradyumn Sharma, Rakesh Vaiwala, Amar Krishna Gopinath, Rajalakshmi Chockalingam, and K Ganapathy Ayappa. Structure of the bacterial cell envelope and interactions with antimicrobials: insights from molecular dynamics simulations. Langmuir, 40(15):7791–7811, 2024.

[4] John-Jackson Yang, Ting-Wei Chang, Yong Jiang, Hsin-Jou Kao, Bin-Hao Chiou, Ming-Shan Kao, and Chun-Ming Huang. Commensal Staphylococcus aureus provokes immunity to protect against skin infection of methicillin-resistant Staphylococcus aureus. Int. J. Mol. Sci., 19(5):1290, 2018.

[5] Leslie Landemaine, Gregory Da Costa, Elsa Fissier, Carine Francis, Stanislas Morand, Jonathan Verbeke, Marie-Laure Michel, Romain Briandet, Harry Sokol, and Audrey Gueniche. Staphylococcus epidermidis isolates from atopic or healthy skin have opposite effect on skin cells: potential implication of the AHR pathway modulation. Front. Immunol., 14:1098160, 2023.

[6] Vishnuvardhan Reddy Sultanpuram, Thirumala Mothe, Sasikala Chintalapati, and Venkata Ramana Chintalapati. Nesterenkonia cremea sp. nov., a bacterium isolated from a soda lake. Int. J. Syst. Evol. Microbiol., 67(6):1861–1866, 2017.

[7] Nina Gunde-Cimerman, Ana Plemenitaš, and Aharon Oren. Strategies of adaptation of microorganisms of the three domains of life to high salt concentrations. FEMS Microbiol. Rev., 42(3):353–375, 2018.

[8] Gwendolyn J Gregory and E Fidelma Boyd. Stressed out: Bacterial response to high salinity using compatible solute biosynthesis and uptake systems, lessons from Vibrionaceae. Comput. Struct. Biotechnol. J., 19:1014–1027, 2021.

[9] Thomas J Piggot, Daniel A Holdbrook, and Syma Khalid. Electroporation of the E. coli and S. aureus membranes: molecular dynamics simulations of complex bacterial membranes. J. Phys.Chem. B, 115(45):13381–13388, 2011.

[10] Sarah Witzke, Michael Petersen, Timothy S Carpenter, and Syma Khalid. Molecular dynamics simulations reveal the conformational flexibility of lipid II and its loose association with the defensin plectasin in the Staphylococcus aureus membrane. Biochemistry, 55(23):3303–3314, 2016.

[11] WF Drew Bennett, Stephen J Fox, Delin Sun, and C Mark Maupin. Bacterial membranes are more perturbed by the asymmetric versus symmetric loading of amphiphilic molecules. Membranes, 12(4):350, 2022.

[12] Faramarz Joodaki, Lenore M Martin, and Michael L Greenfield. Generation and computational characterization of a complex Staphylococcus aureus lipid bilayer. Langmuir, 38(31):9481–9499, 2022.

[13] Davood Zaeifi, Ali Najafi, and Reza Mirnejad. Molecular dynamics simulation of antimicrobial peptide CM15 in Staphylococcus aureus and Escherichia coli model bilayer lipid. Iran. J. Biotechnol., 21(2):e3344, 2023.

[14] Callum Waller, Jan K Marzinek, Eilish McBurnie, Peter J Bond, Philip TF Williamson, and Syma Khalid. Impact on S. aureus and E. coli membranes of treatment with chlorhexidine and alcohol solutions: insights from molecular simulations and nuclear magnetic resonance. J. Mol. Biol., 435(11):167953, 2023.

[15] Barmak Mostofian, Tony Zhuang, Xiaolin Cheng, and Jonathan D Nickels. Branched-chain fatty acid content modulates structure, fluidity, and phase in model microbial cell membranes. J. Phys. Chem. B, 123(27):5814–5821, 2019.

[16] Matthew W Frank, Sarah G Whaley, and Charles O Rock. Branched-chain amino acid metabolism controls membrane phospholipid structure in Staphylococcus aureus. J. Biol. Chem., 297(5), 2021.

[17] Joseph B Lim and Jeffery B Klauda. Lipid chain branching at the iso- and anteiso-positions in complex chlamydia membranes: A molecular dynamics study. Biochimica et Biophysica Acta (BBA) - Biomembranes, 1808(1):323–331, 2011.

[18] Diego de Mendoza and John E Cronan. Thermal regulation of membrane lipid fluidity in bacteria. Trends Biochem. Sci., 8(2):49–52, 1983.

[19] Frederic L Hoch. Cardiolipins and biomembrane function. Biochimica et Biophysica Acta (BBA)- Rev. Biomembranes, 1113(1):71–133, 1992.

[20] Martin Dahlberg. Polymorphic phase behavior of cardiolipin derivatives studied by coarse-grained molecular dynamics. J. Phys. Chem. B, 111(25):7194–7200, 2007.

[21] Martin Dahlberg and Arnold Maliniak. Molecular dynamics simulations of cardiolipin bilayers. J. Phys. Chem. B, 112(37):11655–11663, 2008.

[22] Blake A Wilson, Arvind Ramanathan, and Carlos F Lopez. Cardiolipin-dependent properties of model mitochondrial membranes from molecular simulations. Biophys. J., 117(3):429–444, 2019.

[23] Ruthven NAH Lewis and Ronald N McElhaney. The physicochemical properties of cardiolipin bilayers and cardiolipin-containing lipid membranes. Biochimica et Biophysica Acta (BBA) - Biomembranes, 1788(10):2069–2079, 2009.

[24] Robin A Corey, Wanling Song, Anna L Duncan, T Bertie Ansell, Mark SP Sansom, and Phillip J Stansfeld. Identification and assessment of cardiolipin interactions with E. coli inner membrane proteins. Sci. Adv., 7(34):eabh2217, 2021.

[25] Michał A Surma, Mathias J Gerl, Ronny Herzog, Jussi Helppi, Kai Simons, and Christian Klose. Mouse lipidomics reveals inherent flexibility of a mammalian lipidome. Sci. Rep., 11(1):19364, 2021.

[26] Ronny Herzog, Dominik Schwudke, Kai Schuhmann, Julio L Sampaio, Stefan R Bornstein, Michael Schroeder, and Andrej Shevchenko. A novel informatics concept for high-throughput shotgun lipidomics based on the molecular fragmentation query language. Genome Biol., 12(1):R8, 2011.

[27] Vitor Teixeira, Maria J Feio, and Margarida Bastos. Role of lipids in the interaction of antimicrobial peptides with membranes. Prog. Lipid Res., 51(2):149–177, 2012.

[28] Nermina Malanovic and Karl Lohner. Gram-positive bacterial cell envelopes: The impact on the activity of antimicrobial peptides. Biochimica et Biophysica Acta (BBA)- Biomembranes, 1858(5):936–946, 2016.

[29] Jeffery B Klauda, Richard M Venable, J Alfredo Freites, Joseph W O’Connor, Douglas J Tobias, Carlos Mondragon-Ramirez, Igor Vorobyov, Alexander D MacKerell Jr, and Richard W Pastor. Update of the CHARMM all-atom additive force field for lipids: validation on six lipid types. J. Phys. Chem. B, 114(23):7830–7843, 2010.

[30] RW Pastor and AD MacKerell Jr. Development of the CHARMM force field for lipids. J. Phys. Chem. Lett., 2(13):1526–1532, 2011.

[31] Kenno Vanommeslaeghe, E Prabhu Raman, and Alexander D MacKerell Jr. Automation of the CHARMM General Force Field (CGenFF) II: assignment of bonded parameters and partial atomic charges. J. Chem. Inf. Model., 52(12):3155–3168, 2012.

[32] Leandro Martínez, Ricardo Andrade, Ernesto G Birgin, and José Mario Martínez. PACKMOL: A package for building initial configurations for molecular dynamics simulations. J. Comput. Chem., 30(13):2157–2164, 2009.

[33] Christopher J Knight and Jochen S Hub. MemGen: A general web server for the setup of lipid membrane simulation systems. Bioinformatics, 31(17):2897–2899, 2015.

[34] Mark E Tuckerman. Statistical mechanics: theory and molecular simulation. Oxford university press, 2023.

[35] William Humphrey, Andrew Dalke, and Klaus Schulten. VMD – Visual Molecular Dynamics. J. Mol. Graph., 14:33–38, 1996.

[36] Herman JC Berendsen, JPM van Postma, Wilfred F Van Gunsteren, ARHJ DiNola, and Jan R Haak. Molecular dynamics with coupling to an external bath. J. Chem. Phys., 81(8):3684–3690, 1984.

[37] Glenn J Martyna, Michael L Klein, and Mark Tuckerman. Nosé–Hoover chains: The canonical ensemble via continuous dynamics. J. Chem. Phys., 97(4):2635–2643, 1992.

[38] Michele Parrinello and Aneesur Rahman. Polymorphic transitions in single crystals: A new molecular dynamics method. J. Appl. Phys., 52(12):7182–7190, 1981.

[39] Berk Hess, Henk Bekker, Herman JC Berendsen, and Johannes GEM Fraaije. LINCS: a linear constraint solver for molecular simulations. J. Comput. Chem., 18(12):1463–1472, 1997.

[40] Tom Darden, Darrin York, and Lee Pedersen. Particle mesh Ewald: An Nlog(N) method for Ewald sums in large systems. J. Chem. Phys., 98(12):10089–10092, 1993.

[41] Giovanni Bussi, Davide Donadio, and Michele Parrinello. Canonical sampling through velocity rescaling. J. Chem. Phys., 126(1):014101, 2007.

[42] Ilario G Tironi, René Sperb, Paul E Smith, and Wilfred F van Gunsteren. A generalized reaction field method for molecular dynamics simulations. J. Chem. Phys., 102(13):5451–5459, 1995.

[43] Alžbeta Kubincová, Sereina Riniker, and Philippe H Hünenberger. Reaction-field electrostatics in molecular dynamics simulations: Development of a conservative scheme compatible with an atomic cutoff. Phys. Chem. Chem. Phys., 22(45):26419–26437, 2020.

[44] Rakesh Vaiwala and K Ganapathy Ayappa. Martini-3 coarse-grained models for the bacterial lipopolysaccharide outer membrane of Escherichia coli. J. Chem. Theory Comput., 20(4):1704–1716, 2023.

[45] Thomas J Piggot, Jane R Allison, Richard B Sessions, and Jonathan W Essex. On the calculation of acyl chain order parameters from lipid simulations. J. Chem. Theory Comput., 13(11):5683–5696, 2017.

[46] Etana Padan, Eitan Bibi, Masahiro Ito, and Terry A Krulwich. Alkaline pH homeostasis in bacteria: new insights. Biochimica et biophysica acta (BBA) - Biomembranes, 1717(2):67–88, 2005.

[47] Elena Beltrán-Heredia, Feng-Ching Tsai, Samuel Salinas-Almaguer, Francisco J Cao, Patricia Bassereau, and Francisco Monroy. Membrane curvature induces cardiolipin sorting. Commun. Biol., 2(1):225, 2019.

[48] Alex H De Vries, Alan E Mark, and Siewert J Marrink. The binary mixing behavior of phospholipids in a bilayer: a molecular dynamics study. J. Phys. Chem. B, 108(7):2454–2463, 2004.

[49] Siewert J Marrink, Alex H De Vries, and Alan E Mark. Coarse grained model for semiquantitative lipid simulations. J. Phys. Chem. B, 108(2):750–760, 2004.

[50] James R Sheats and Harden M McConnell. A photochemical technique for measuring lateral diffusion of spin-labeled phospholipids in membranes. Proc. Natl. Acad. Sci., 75(10):4661–4663, 1978.

[51] An-Li Kuo and Charles G Wade. Lipid lateral diffusion by pulsed nuclear magnetic resonance. Biochemistry, 18(11):2300–2308, 1979.

[52] Wolfgang Helfrich. Out-of-plane fluctuations of lipid bilayers. Z. Naturforsch. C, 30(11-12):841–842, 1975.

[53] Wolfgang Helfrich. Steric interaction of fluid membranes in multilayer systems. Z. Naturforsch. A, 33(3):305–315, 1978.

[54] Rakesh Vaiwala and Rochish Thaokar. Probing entropic repulsion through mesoscopic simulations. Europhys. Lett., 120(4):48001, 2018.

[55] Foram M Thakkar, Prabal K Maiti, V Kumaran, and KG Ayappa. Verifying scalings for bending rigidity of bilayer membranes using mesoscale models. Soft Matter, 7(8):3963–3966, 2011.

[56] Erik G Brandt, Anthony R Braun, Jonathan N Sachs, John F Nagle, and Olle Edholm. Interpretation of fluctuation spectra in lipid bilayer simulations. Biophys. J., 100(9):2104–2111, 2011.

[57] Sam Brown, Jessica Pallarez, and Marat R Talipov. Comparative analysis of bending moduli in one-component membranes via coarse-grained molecular dynamics simulations. Biophys. J., 2025.

[58] Lars D Renner and Douglas B Weibel. Cardiolipin microdomains localize to negatively curved regions of Escherichia coli membranes. Proc. Natl. Acad. Sci., 108(15):6264–6269, 2011.

[59] Tatyana Romantsov, Ziqiang Guan, and Janet M Wood. Cardiolipin and the osmotic stress responses of bacteria. Biochimica et Biophysica Acta (BBA) - Biomembranes, 1788(10):2092–2100, 2009.

[60] Georg Pabst, Aden Hodzic, Janez Štrancar, Sabine Danner, Michael Rappolt, and Peter Laggner. Rigidification of neutral lipid bilayers in the presence of salts. Biophys. J., 93(8):2688–2696, 2007.

[61] Judith U De Mel, Sudipta Gupta, Rasangi M Perera, Ly Ngo, Piotr Zolnierczuk, Markus Bleuel, Sai Venkatesh Pingali, and Gerald J Schneider. Influence of external NaCl salt on membrane rigidity of neutral DOPC vesicles. Langmuir, 36(32):9356–9367, 2020.

[62] MMAE Claessens, BF Van Oort, FAM Leermakers, FA Hoekstra, and MA Cohen Stuart. Charged lipid vesicles: effects of salts on bending rigidity, stability, and size. Biophys. J., 87(6):3882–3893, 2004.

[63] Teshani Kumarage, Sudipta Gupta, Nicholas B Morris, Fathima T Doole, Haden L Scott, Laura-Roxana Stingaciu, Sai Venkatesh Pingali, John Katsaras, George Khelashvili, Milka Doktorova, Michael F Brown, and Rana Ashkar. Cholesterol modulates membrane elasticity via unified biophysical laws. Nature Commun., 16(1):7024, 2025.

[64] Pablo Campomanes, Valeria Zoni, and Stefano Vanni. Local accumulation of diacylglycerol alters membrane properties nonlinearly due to its transbilayer activity. Commun. Chem., 2(1):72, 2019.

[65] RJ Bruckner, Sheref Samir Mansy, A Ricardo, L Mahadevan, and JW Szostak. Flip-flop-induced relaxation of bending energy: implications for membrane remodeling. Biophys. J., 97(12):3113–3122, 2009.

[66] Rakesh Vaiwala, Pradyumn Sharma, Mrinalini Puranik, and K Ganapathy Ayappa. Developing a coarse-grained model for bacterial cell walls: evaluating mechanical properties and free energy barriers. J. Chem. Theory Comput., 16(8):5369–5384, 2020.

[67] Rakesh Vaiwala, Pradyumn Sharma, and K Ganapathy Ayappa. Differentiating interactions of antimicrobials with Gram-negative and Gram-positive bacterial cell walls using molecular dynamics simulations. Biointerphases, 17(6):061008, 2022.

[68] T Eklund. Inhibition of growth and uptake processes in bacteria by some chemical food preservatives. J. Appl. Bacteriol., 48(3):423–432, 1980.

[69] JD Baranowski and CW Nagel. Antimicrobial and antioxidant activities of alkyl hydroxycinnamates (alkacins) in model systems and food products. Can. Inst. Food Sci. Technol. J., 17(2):79–85, 1984.

[70] Jochen Weiss, Myriam Loeffler, and Nino Terjung. The antimicrobial paradox: why preservatives lose activity in foods. Curr. Opin. Food Sci., 4:69–75, 2015.

[71] Rajalakshmi Chockalingam, Rakesh Vaiwala, Nivedita Patil, and K Ganapathy Ayappa. Preservative kinetics and influence of surfactants on Gram-negative bacterial membrane partitioning. Langmuir, 2026.

[72] R Becerril, S Manso, C Nerin, and R Gómez-Lus. Antimicrobial activity of Lauroyl Arginate Ethyl (LAE), against selected food-borne bacteria. Food Control, 32(2):404–408, 2013.

[73] Pradyumn Sharma and K Ganapathy Ayappa. A molecular dynamics study of antimicrobial peptide interactions with the lipopolysaccharides of the outer bacterial membrane. J. Membr. Biol., pages 1–11, 2022.

[74] M Moosazadeh Moghaddam, F Abolhassani, H Babavalian, R Mirnejad, K Azizi Barjini, and J Amani. Comparison of in vitro antibacterial activities of two cationic peptides CM15 and CM11 against five pathogenic bacteria: Pseudomonas aeruginosa, Staphylococcus aureus, Vibrio cholerae, Acinetobacter baumannii, and Escherichia coli. Probiotics Antimicrob. Proteins, 4(2):133–139, 2012.

[75] Avijeet Kulshrestha, Nishant Bahuguna, Sudeep Punnathanam, and Ganapathy Ayappa. BPS2025-cardiolipin in the bacterial membranes determines the binding of the CM15 antimicrobial peptide. Biophys. J., 124(3):187a, 2025.

[76] Sara Pistolesi, Rebecca Pogni, and Jimmy B Feix. Membrane insertion and bilayer perturbation by antimicrobial peptide CM15. Biophys. J., 93(5):1651–1660, 2007.

[77] Pradyumn Sharma, Srividhya Parthasarathi, Nivedita Patil, Morris Waskar, Janhavi S Raut, Mrinalini Puranik, K Ganapathy Ayappa, and Jaydeep Kumar Basu. Assessing barriers for antimicrobial penetration in complex asymmetric bacterial membranes: a case study with thymol. Langmuir, 36(30):8800–8814, 2020.

[78] Rakesh Gupta, Beena Rai, and K Ganapathy Ayappa. Interaction of Choline-Based Ionic Liquids with Model Lipid Membranes: Force-Field Parametrization and Membrane Partitioning. J. Phys. Chem. B, 130(12):3342–3357, 2026.

[79] Hiroshi Onishi and Masahiro Kamekura. Micrococcus halobius sp. n. Int. J. Syst. Evol. Microbiol., 22(4):233–236, 1972.

[80] Matthew D Collins, Paul A Lawson, Matthias Labrenz, Brian J Tindall, N Weiss, and Peter Hirsch. Nesterenkonia lacusekhoensis sp. nov., isolated from hypersaline Ekho Lake, East Antarctica, and emended description of the genus Nesterenkonia. Int. J. Syst. Evol. Microbiol., 52(4):1145–1150, 2002.

