## Supporting Information for "Molecular models for Gram-positive bacterial strains: Assessing membrane properties and small molecule interactions for *S. aureus*, *S. epidermidis* and *N. lacusekhoensis*"

Rakesh Vaiwala [1], Ernest Christy [2], Morris Waskar [2] and K. Ganapathy Ayappa [1,*]

[1] *Department of Chemical Engineering, Indian Institute of Science, Bangalore 560012, India*
[2] *Unilever Research and Development, Bangalore 560066, India*
*

**Table S1:** Relative abundances of major fatty acyl chains detected in *S. epidermidis* ATCC 12228 and *N. lacusekhoensis*, highlighting the predominance of iso and anteiso branched fatty acids. Data represent the proportional distribution of quantified fatty acids as determined by shotgun lipidomics.

| **S. epidermidis** | | **N. lacusekhoensis** | |
|---|---|---|---|
| **Fatty acyl chain** | **Amount in %** | **Fatty acyl chain** | **Amount in %** |
| FA 15:0 (anteiso) | 31.49 | FA 16:0 | 32.17 |
| FA 15:0 (iso) | 14.04 | FA 18:1 | 23.50 |
| FA 20:0 | 13.04 | FA 15:O (anteiso) | 13.82 |
| FA 14:0 (iso) | 10.64 | FA 16:0 (iso) | 11.69 |
| FA 18:0 | 7.42 | FA 17:0 (anteiso) | 9.16 |
| Other FA chains | 23.36 | Other FA chains | 9.66 |

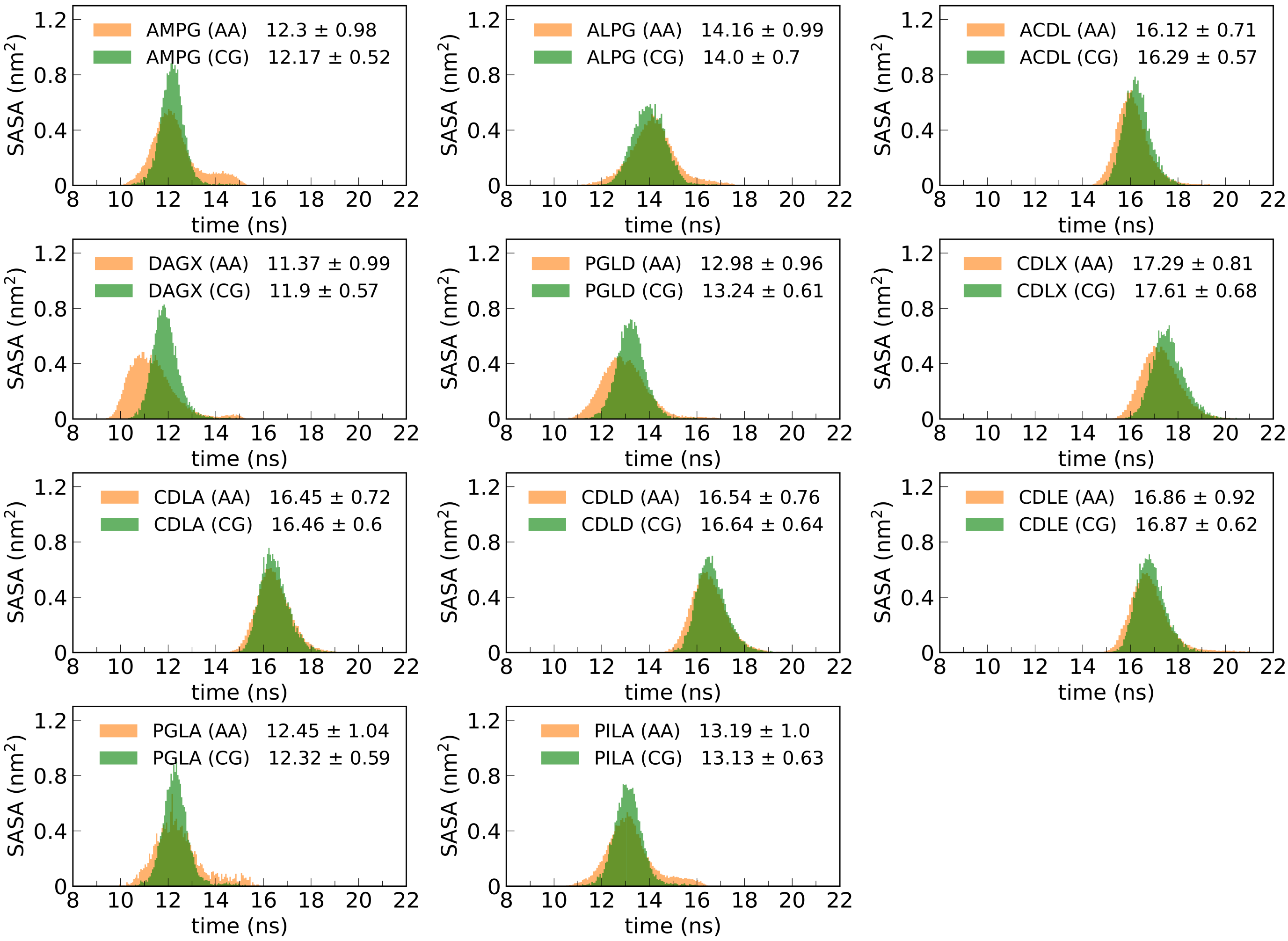


**Figure S2:** The histograms for solvent accessible surface area (SASA) of individual lipid models. Each histogram panel corresponds to all-atom (AA) and coarse-grained (CG) simulations of a single lipid molecule in bulk water.

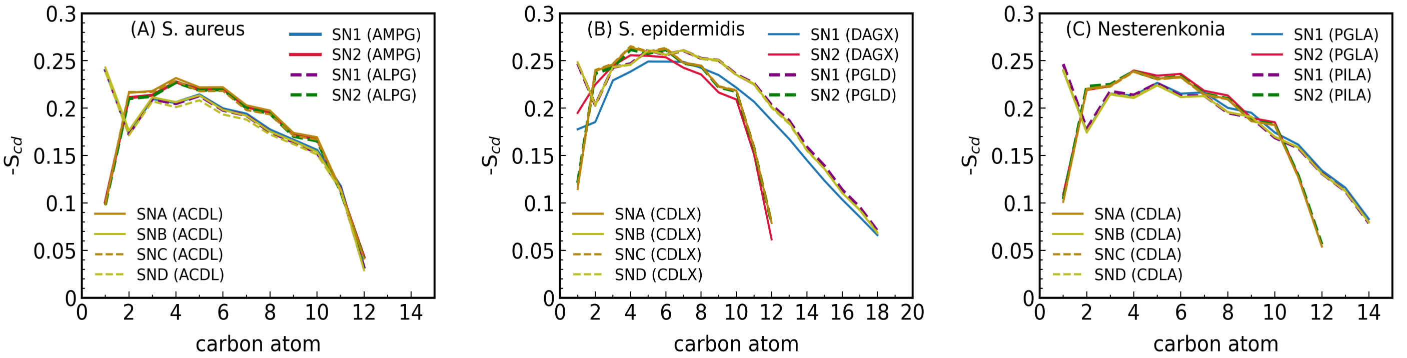


**Figure S3:** Acyl tail order parameters for individual lipids from all-atom simulations of the membranes for different Gram-positive bacteria strains.

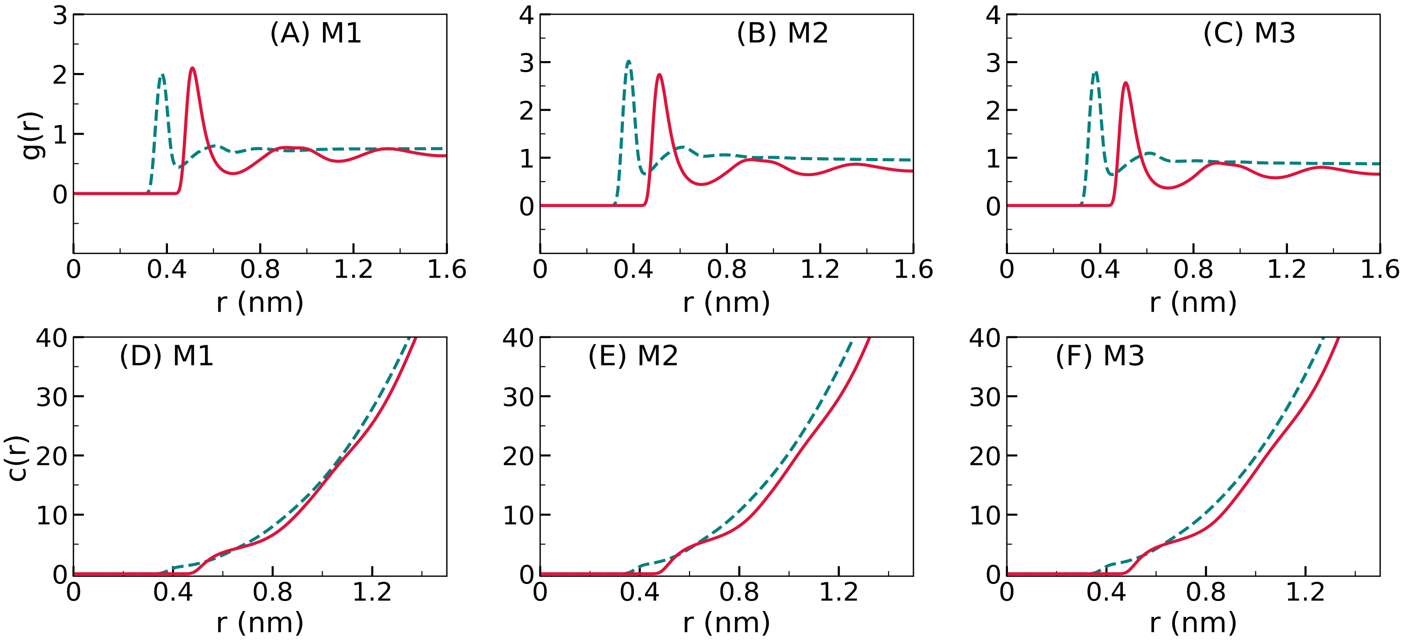


**Figure S4:** (A-C) Radial distribution functions for ordering of water around phosphorus atoms in AA (phosphate beads in CG) in *S. aureus* (M1), *S. epidermidis* (M2) and *N. lacusekhoensis* (M3) membranes. (D-F) The coordination numbers for water around the phosphorus atoms in AA and phosphate beads in CG membranes, indicating hydration of lipid headgroups. The dotted lines refer to atomistic simulation data, and the solid lines represent the coarse-grained simulations. The coordination number for water molecules computed from atomistic simulations is scaled down by factor 4 to compare it with the coarse-grained water beads according to 4:1 mapping scheme.

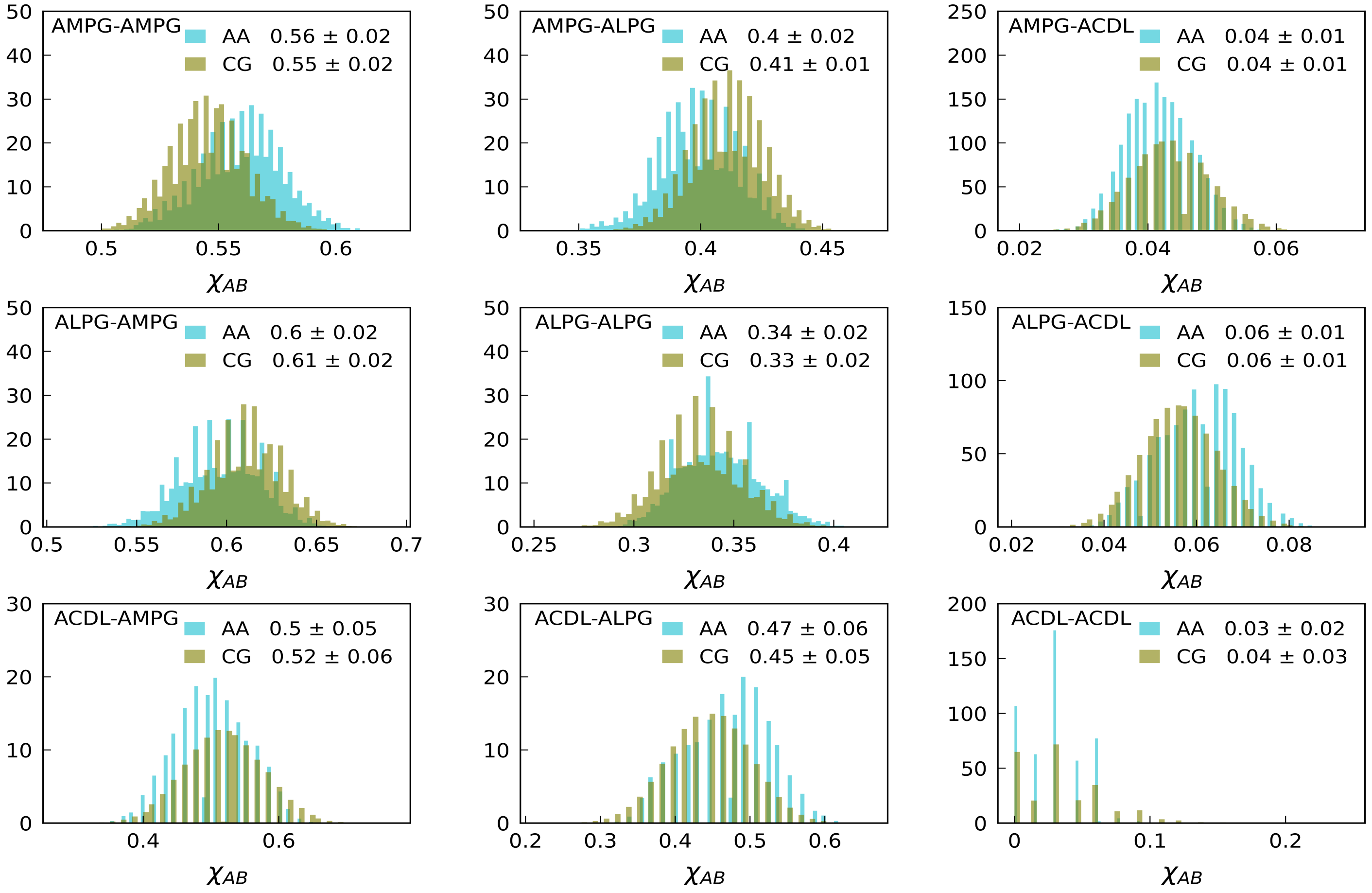


**Figure S5:** The histograms showing fractions of nearest neighbors for each type of lipid in *S. aureus* membrane simulations. The mean and standard deviation are indicated in legends.

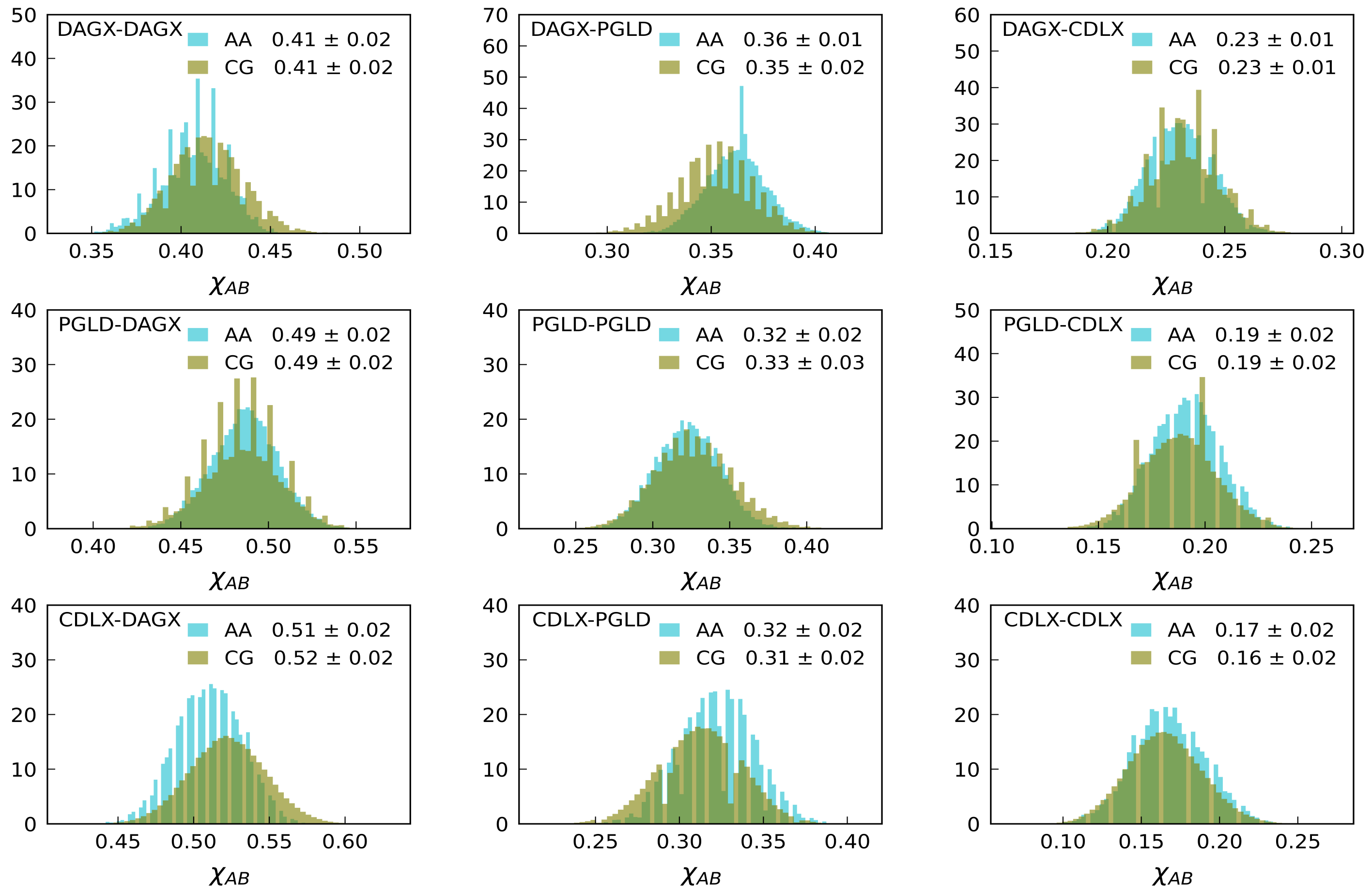


**Figure S6:** The histograms showing fractions of nearest neighbors for each type of lipid in *S. epidermidis* membrane simulations. The statistics for mean and standard deviation are indicated in legends.

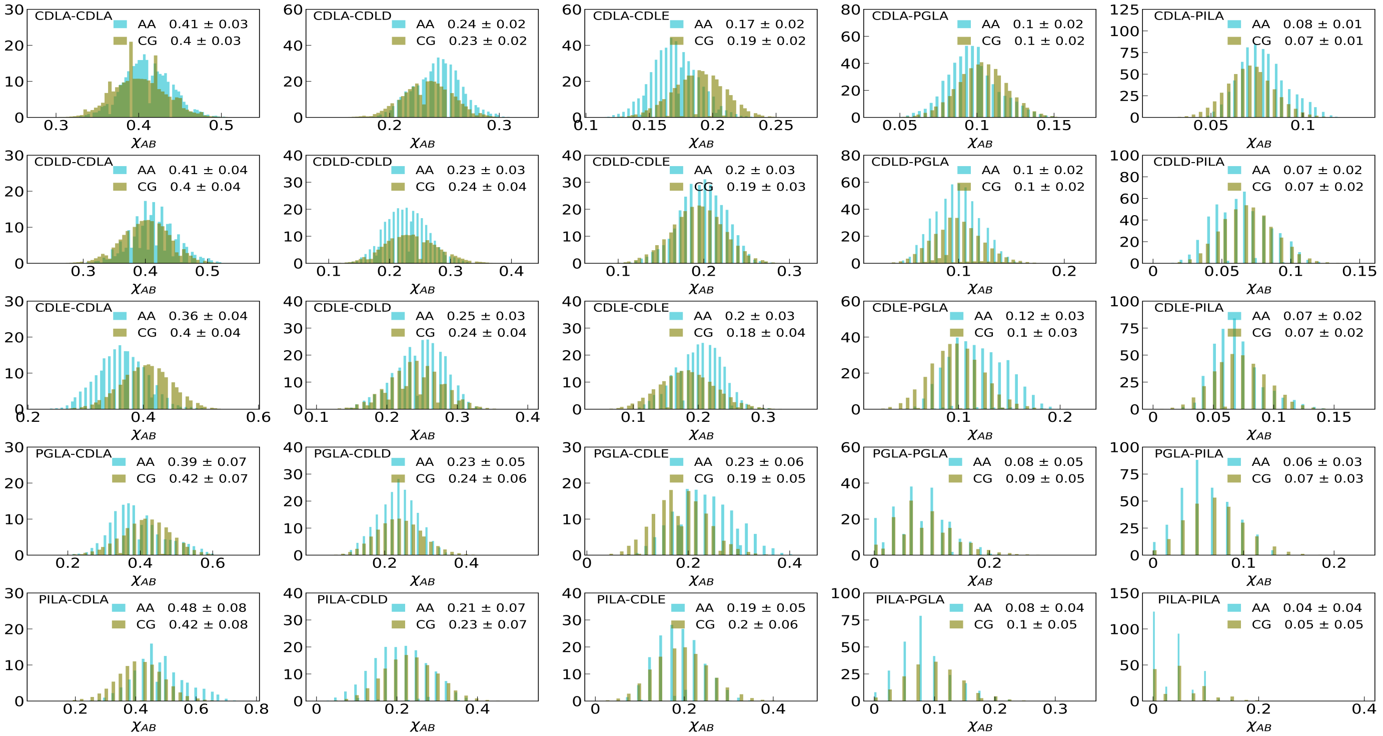


**Figure S7:** The histograms showing fractions of nearest neighbors for each type of lipid in *N. lacusekhoensis* membrane simulations. The data for mean and standard deviation are provided in legends.

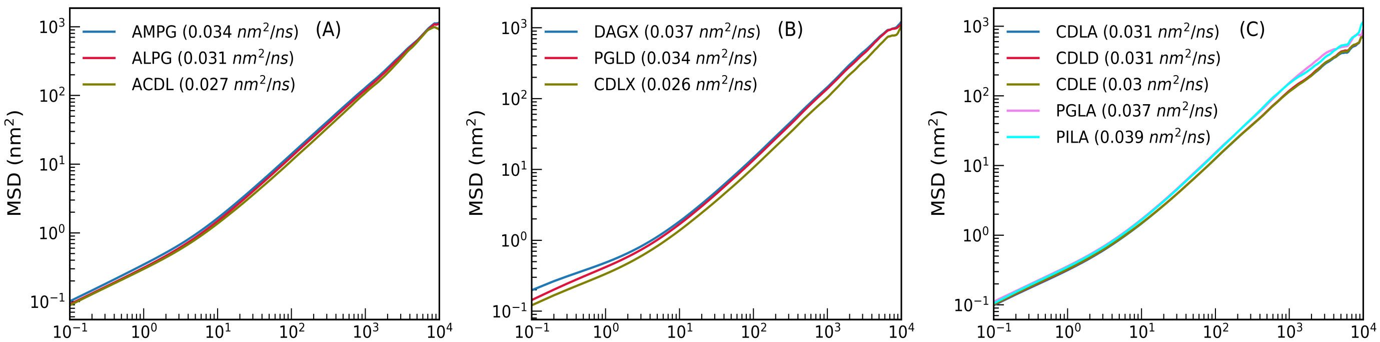


**Figure S8:** The mean square displacements (MSD) for CG membrane lipids in (A) *S. aureus* (B) *S. epidermidis*, (C) *N. lacusekhoensis* membranes.

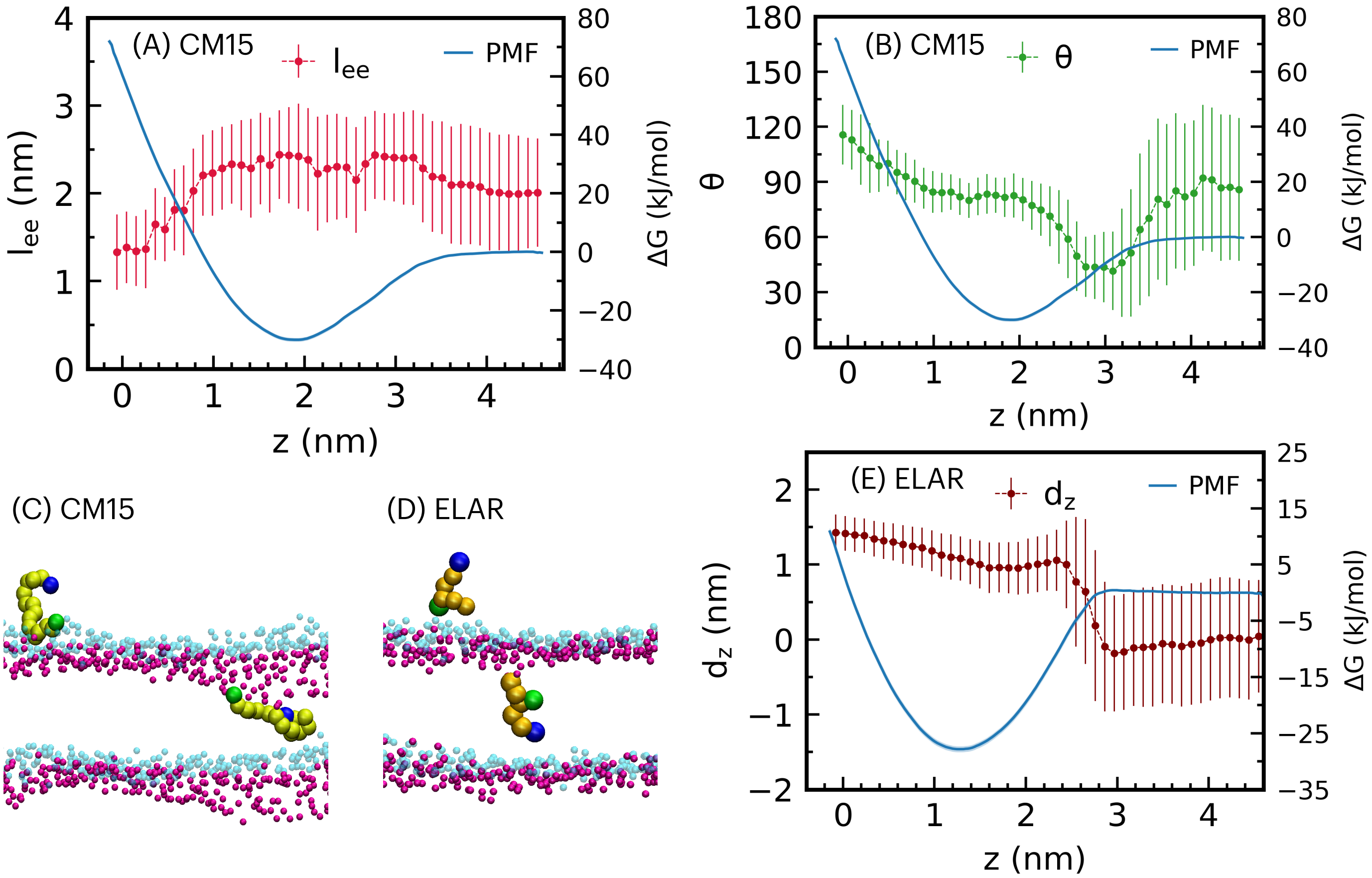


**Figure S9:** (A-C) Conformational changes of CM15 peptide interacting with *S. aureus* CG membrane. (A) Peptide end-to-end distance ($l_{ee}$) as peptide approaches the membrane center. (B) The angle ($\theta$) variation with distance z. The angle is defined between the vector connecting terminal backbone beads and the membrane normal (z-axis). (C-D) The simulation snapshots were taken at instances when the molecule was outside the membrane (headgroups in translucent phosphate beads shown in cyan color) and when it was at membrane center (phosphate headgroup beads in crimson color). The peptide backbone beads are indicated by lime color with N- and C-terminal beads shown in green and blue colors, respectively. The charged amine group and terminal carbon bead of fatty chain of ELAR are shown in green and blue colors, repsectively. (E) The preferential orientation of ELAR interacting with CG membrane of *S. aureus*. The y-axis label $d_z$ is the z-component of the distance between the charged amine bead and the terminal carbon bead. In above panels, the free energy ($\Delta G$) is also indicated as a function of molecule's distance from the membrane center (z), and the error bars are the standard deviations of the data.

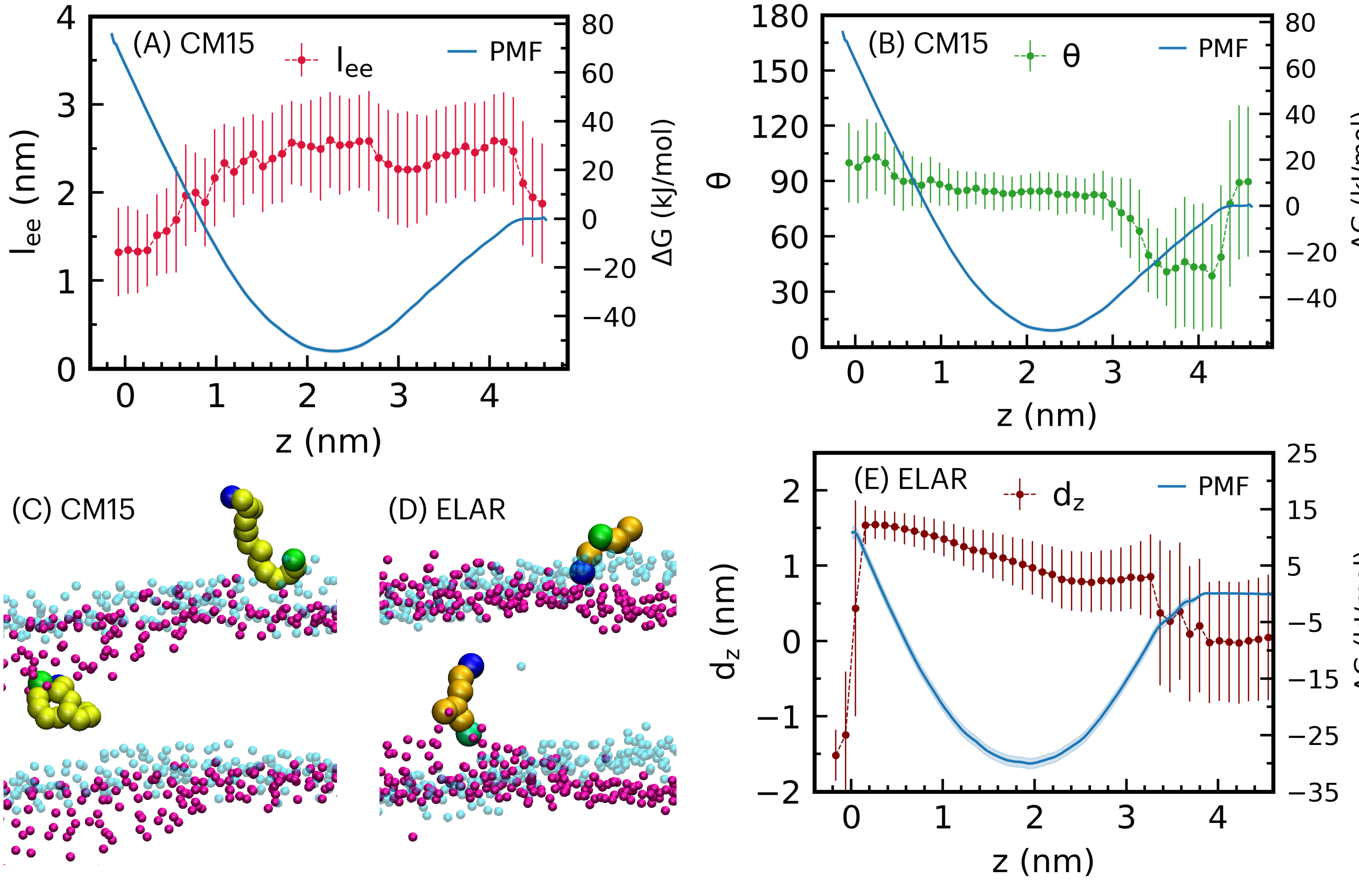


**Figure S10:** (A-C) Conformational changes of CM15 peptide interacting with *S. epidermidis* CG membrane. (A) Peptide end-to-end distance ($l_{ee}$) as peptide approaches the membrane center. (B) The angle ($\theta$) variation with distance z. The angle is defined between the vector connecting terminal backbone beads and the membrane normal (z-axis). (C-D) The simulation snapshots were taken at instances when the molecule was outside the membrane (headgroups in translucent phosphate beads shown in cyan color) and when it was at membrane center (phosphate headgroup beads in crimson color). The peptide backbone beads are indicated by lime color with N- and C-terminal beads shown in green and blue colors, respectively. The charged amine bead and terminal carbon bead of fatty chain of ELAR (Figure 10 in main text) are shown in green and blue colors, repsectively. (E) The preferential orientation of ELAR interacting with CG membrane of *S. epidermidis*. The y-axis label $d_z$ is the z-component of the distance between the charged amine bead and terminal carbon bead. In above panels, the free energy ($\Delta G$) is also indicated as a function of molecule's distance from the membrane center (z), and the error bars are the standard deviations of the data.

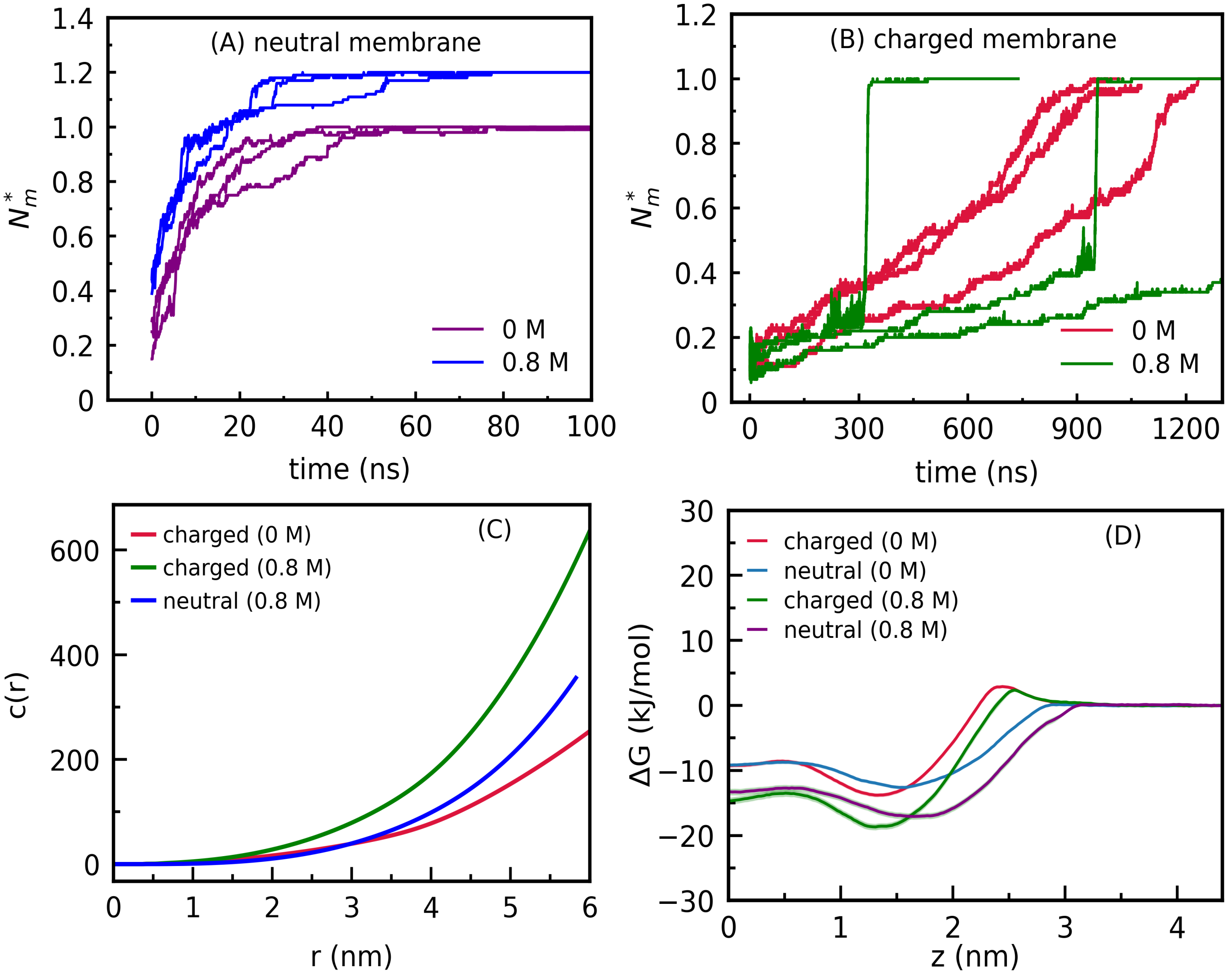


**Figure S11:** (A-B) Fractions ($N_m^*$) of thymol molecules in the neutral and charged atomistic membranes. (C) The coordination numbers for potassium ions around phosphorus atoms for atomistic simulations of thymol in charged and neutral membranes of *N. lacusekhoensis*. (D) Potential of mean force between thymol and charged and neutral coarse-grained membranes of *N. lacusekhoensis* at 0 and 0.8 M salt concentrations.

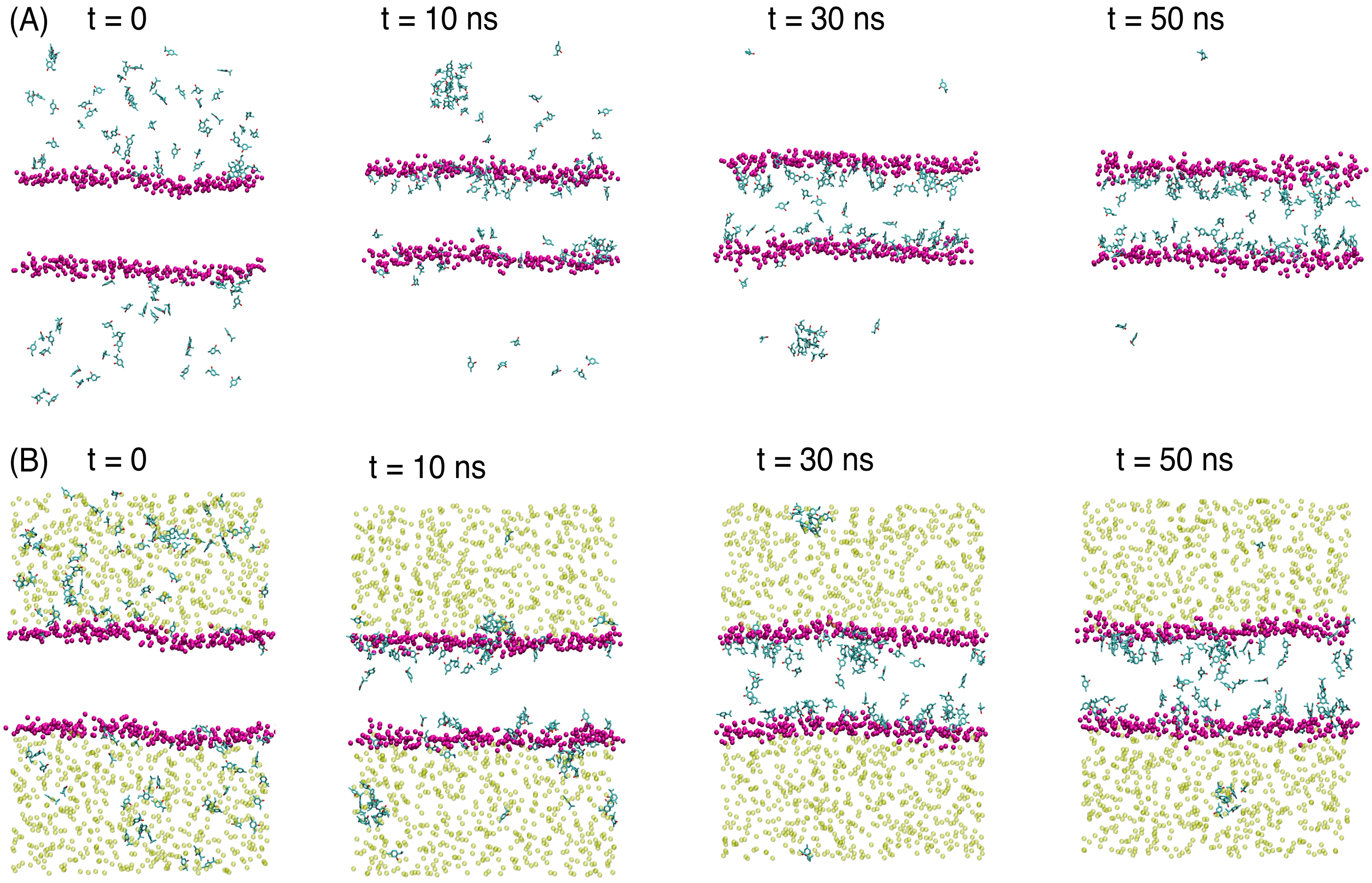


**Figure S12:** The simulation snapshots showing spontaneous uptake of thymol molecules in the atomistic neutral membrane of *N. lakusekhoensis* at (A) 0 M and (B) 0.8 M salt concentrations. The phosphate headgroups are indicated by crimson VDW spheres, potassium ions are shown in translucent VDW spheres, and thymol molecules are depicted in Licorice representation. Water, chloride ions and acyl chains of lipids are not shown for clarity.

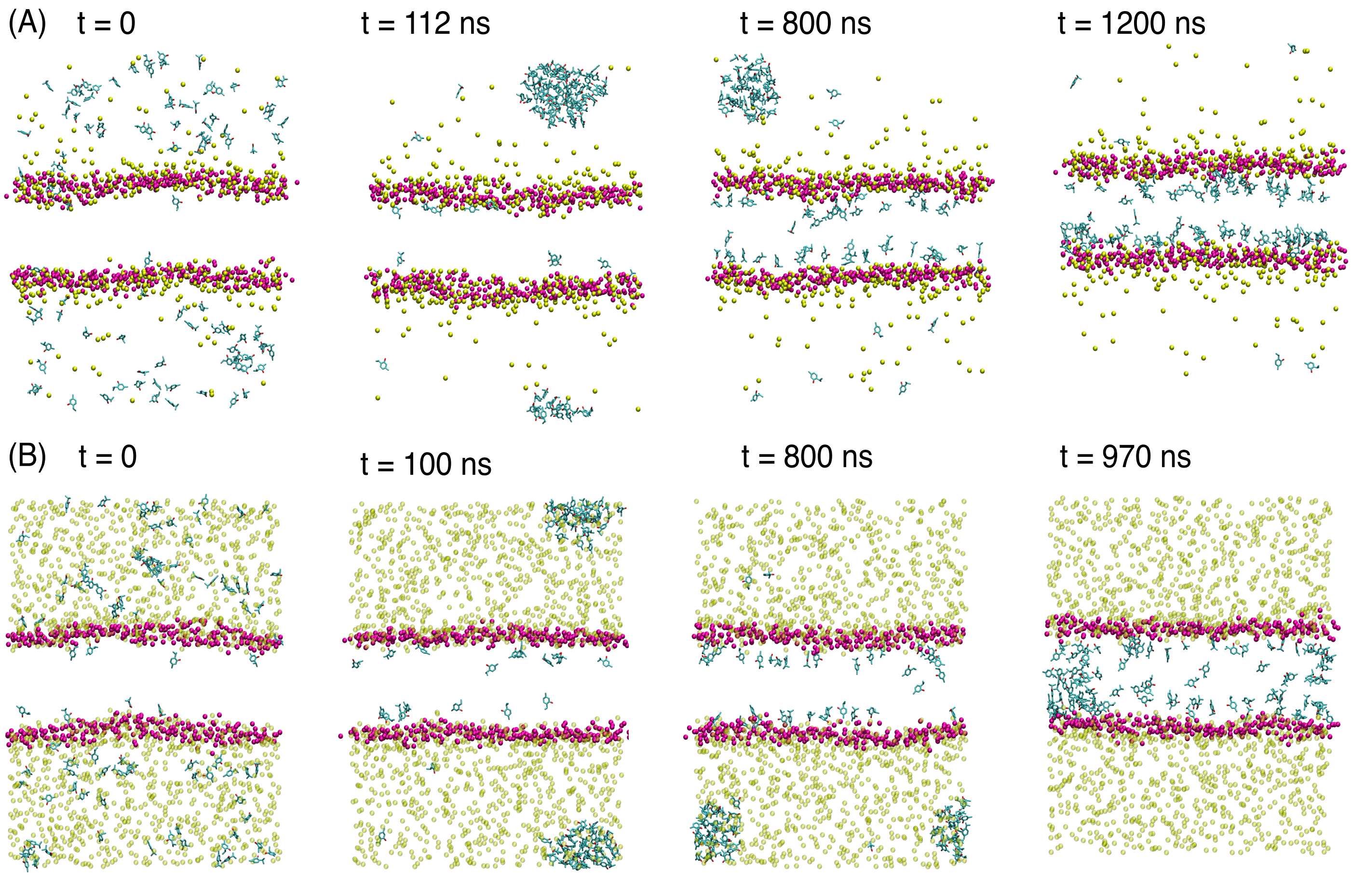


**Figure S13:** The insertion of thymol molecules into the atomistic charged membrane of *N. lakusekhoensis* at (A) 0 M and (B) 0.8 M salt concentrations. The phosphate headgroups are indicated by crimson VDW spheres, potassium ions are shown in translucent VDW spheres, and thymol molecules are depicted in Licorice representation. Water, chloride ions and acyl chains of lipids are not shown for clarity.

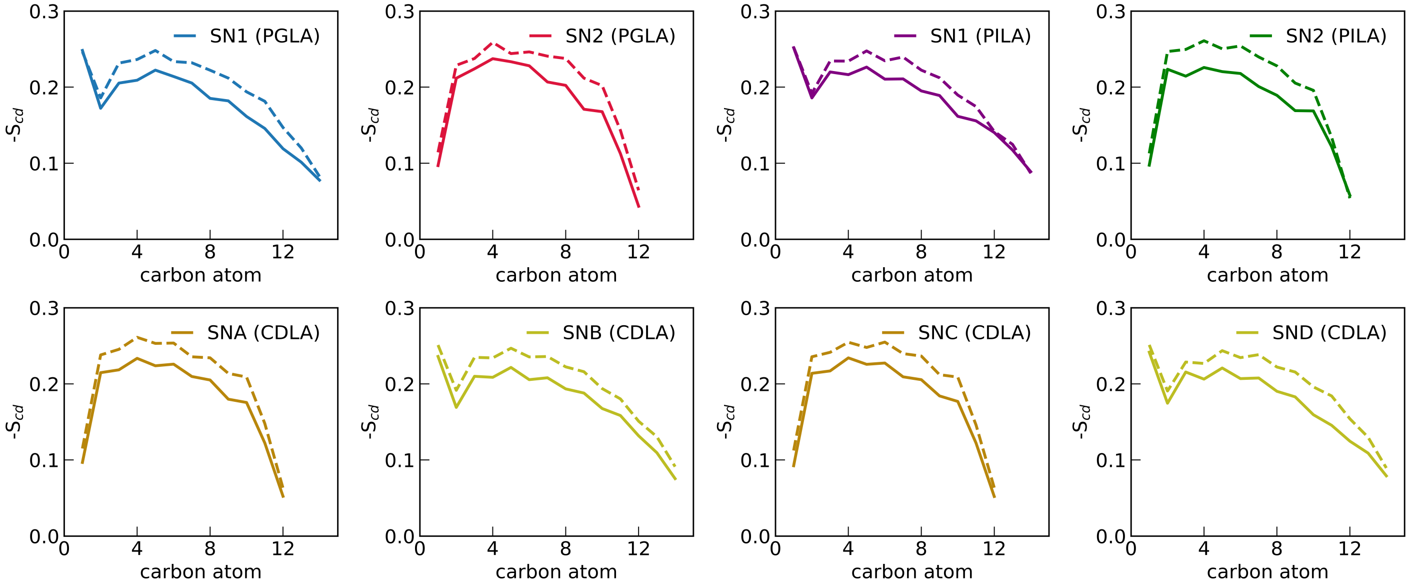


**Figure S14:** The chain order parameters for CDLA, PGLA and PILA lipids in the atomistic charged membrane of *N. lakusekhoensis* at 0 M (solid lines) and 0.8 M (dashed lines) salt concentrations.
